# Vole genomics links determinate and indeterminate growth of teeth

**DOI:** 10.1101/2023.12.18.572015

**Authors:** Zachary T. Calamari, Andrew Song, Emily Cohen, Muspika Akter, Rishi Das Roy, Outi Hallikas, Mona M. Christensen, Pengyang Li, Pauline Marangoni, Jukka Jernvall, Ophir D. Klein

## Abstract

Continuously growing teeth are an important innovation in mammalian evolution, yet genetic regulation of continuous growth by stem cells remains incompletely understood. Dental stem cells responsible for tooth crown growth are lost at the onset of tooth root formation. Genetic signaling that initiates this loss is difficult to study with the ever-growing incisor and rooted molars of mice, the most common mammalian dental model species, because signals for root formation overlap with signals that pattern tooth size and shape (i.e., cusp patterns). Different species of voles (Cricetidae, Rodentia, Glires) have evolved rooted and unrooted molars that have similar size and shape, providing alternative models for studying roots. We assembled a *de novo* genome of *Myodes glareolus*, a vole with high-crowned, rooted molars, and performed genomic and transcriptomic analyses in a broad phylogenetic context of Glires (rodents and lagomorphs) to assess differential selection and evolution in tooth forming genes. We identified 15 dental genes with changing synteny relationships and six dental genes undergoing positive selection across Glires, two of which were undergoing positive selection in species with unrooted molars, *Dspp* and *Aqp1*. Decreased expression of both genes in prairie voles with unrooted molars compared to bank voles supports the presence of positive selection and may underlie differences in root formation. Bulk transcriptomics analyses of embryonic molar development in bank voles also demonstrated conserved patterns of dental gene expression compared to mice, with species-specific variation likely related to developmental timing and morphological differences between mouse and vole molars. Our results support ongoing evolution of dental genes across Glires, revealing the complex evolutionary background of convergent evolution for ever-growing molars.

## INTRODUCTION

Hypselodonty, or the presence of unrooted and thus ever-growing teeth, has evolved multiple times in mammals. Glires—the clade containing rodents, rabbits, and their relatives— have hypselodont incisors (1), and multiple Glires have also evolved hypselodont molars (Fig. 1). At least in rodents, molar hypselodonty evolved considerably later than hypsodont molars, which are high crowned but rooted, which in turn evolved later than hypselodont incisors. In Glires, molars appear to increase in crown height from brachydonty (low-crowned, rooted), through hypsodonty (high-crowned, rooted), toward hypselodonty (high-crowned, unrooted) (2). Mice (*Mus musculus*), the primary mammalian model species of dental research, have hypselodont incisors but retain brachydont molars. Because of this, mice cannot provide information about the hypsodont teeth that likely preceded hypselodonty.

**Figure 1.**
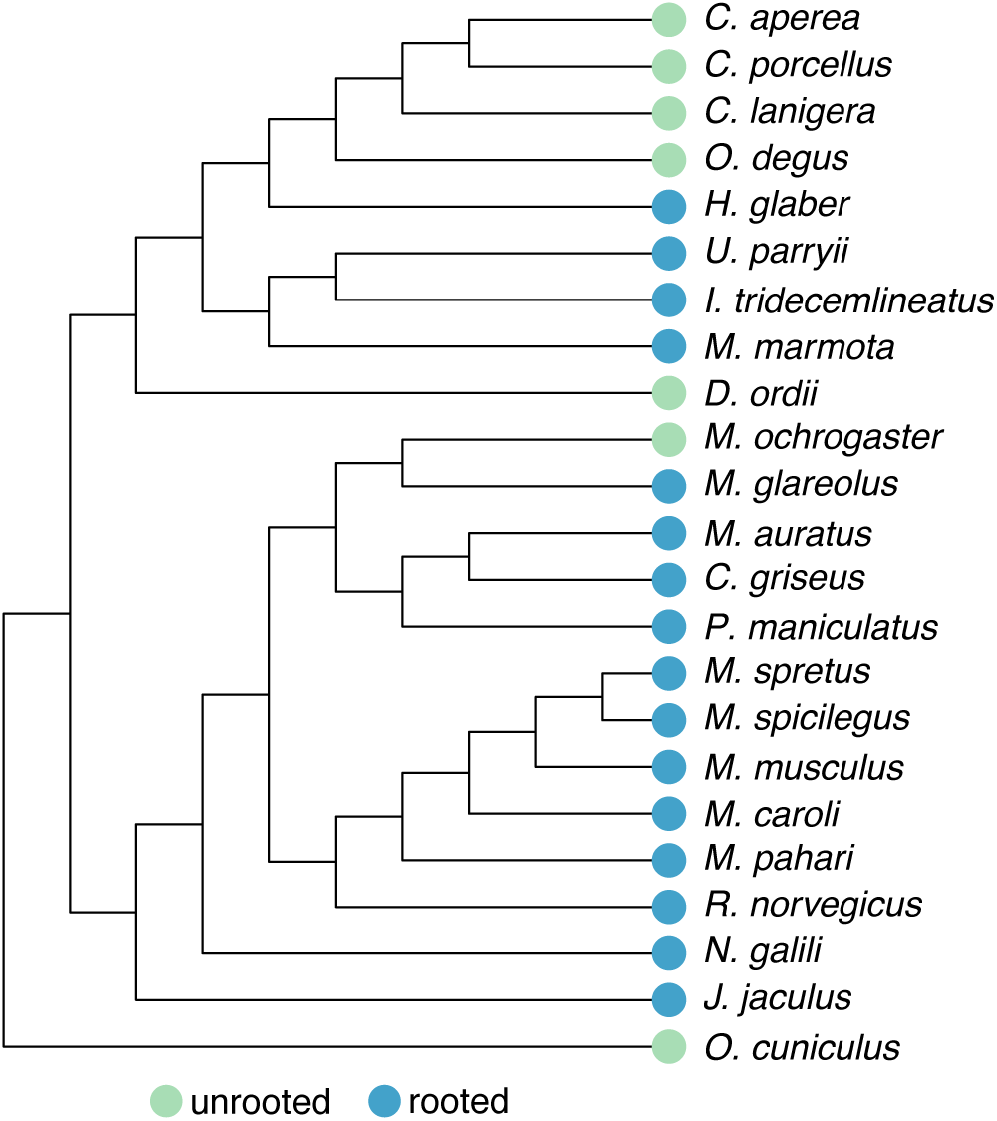
Species tree of Glires based on the Ensembl Compara species tree. Whether each species has rooted or unrooted molars is indicated by colored circles at the tip of each branch. Note that unrooted, or hypselodont, molars have evolved multiple times across Glires. This topology was the basis for our orthology analysis.

Mammalian teeth sit in bony sockets, held in place by soft tissue (periodontal ligament) attached to cementum-covered tooth roots (3). Ligamentous tooth attachment may have arisen along with a reduction in the rate of tooth replacements, providing greater flexibility for repositioning the teeth as the dentary grows (4,3). Consequently, the limited replacement of mammalian teeth (two sets of teeth in most mammals and one in Glires) may have spurred the evolution of hypsodont and hypselodont teeth, both with high crowns that compensate for tooth wear from gritty or phytolith-heavy diets (5,6), and resulted in further modification of the anchoring roots. The convergent evolution of unrooted molars in Glires presents an opportunity to identify whether consistent developmental and genomic changes underlie the formation of hypselodont teeth in different species, in turn revealing the conserved mechanisms that produce tooth roots. Furthermore, the relatively recent evolution of molar hypselodonty, starting in the Middle Miocene (approximately 16-12 Ma) (2), should provide molecular evidence for the steps required to make a continuously growing organ.

Dental development proceeds from the tooth germ, composed of epithelium and mesenchyme, through phases known as the bud, cap, and bell (7). Multipotent enamel epithelium differentiates into the cells that form the tooth crown (8–11). As development progresses in rooted teeth, the epithelium at the tooth apex transitions first to a tissue called Hertwig’s epithelial root sheath, and eventually cementum-covered roots (9,10). Studies have identified numerous candidate genes and pathways with various roles during root development, such as *Fgf10*, which decreases in expression at the beginning of root formation (12–18). Although research on mouse molars has identified genetic signals related to root formation, a number of the key genes studied have broad developmental roles, such as *Wnt* family members (14), or overlap considerably with genes also involved in patterning the size and shape of the tooth (17,19–22). This overlap between shape and root expression patterns confounds our ability to identify a clear signal initiating root formation.

Evolutionary novelties such as high-crowned hypsodont and hypselodont molars can arise from differences in gene expression and regulation (23–26). Evolutionarily conserved gene expression levels produce conserved phenotypes, and changes in gene regulatory networks have long been linked to morphological evolution (27,28). The order of genes along a chromosome (synteny) can affect gene expression and regulation, as regulatory sequences are often located near their target genes (cis-regulatory elements) (29–31). Genome rearrangements that place genes near new regulatory elements may change the expression and selective environment of those genes; these small-scale rearrangements of genes may be common in mammals (32–34). Likewise, regions of chromosomes that form topologically associated domains may experience similar selective pressures, including selection against rearrangement (35,36). Genes involved in molar development are not syntenic in the mouse genome nor are genes with organ-specific expression (37), and thus the regulatory or selection effects of co-localization need not apply to all dental genes at once. Changes in genome architecture between Glires species thus may result in different selective and expression environments for dental genes that could result in the evolution of hypselodont molars.

To establish a model rodent species with hypsodont molars for close comparison to hypselodont molars, we sequenced and annotated a highly-complete *de novo* genome of *Myodes glareolus*, the bank vole. The bank vole is increasingly used in medical and environmental research, ranging from studying zoonotic diseases (38) to immune responses (39,40), and even assessing environmental remediation efforts through heavy metals that accumulate in vole teeth (41,42), thus our efforts may be of use beyond dental research. The bank vole’s hypsodont molars bridge the gap between low-crowned mouse and hypselodont prairie vole (*Microtus ochrogaster*) molars, reducing the effects of morphological differences on root formation signaling. We performed a suite of genomic and transcriptomic tests of our new bank vole genome in a broad phylogenetic context to test the hypothesis that dental genes are undergoing positive selection and exhibit different expression patterns in species with unrooted, hypselodont molars. We predicted that genes without conserved syntenic relationships in these species would be more likely to have sites under positive selection or significantly different expression. Our analyses revealed loss of synteny and positive selection for dental genes in Glires with unrooted molars compared to those with rooted molars. We also demonstrated strong conservation of dental gene expression patterns between bank voles and mice, with key differences related to the timing and patterning of tooth morphology.

## RESULTS

### Orthology assessment and loss of synteny

To identify which sequences in our bank vole (*Myodes glareolus*) genome and annotation had the same evolutionary history as dental genes identified in other Glires and assess genome rearrangements, we performed orthology and synteny analyses in a broad phylogenetic context. OrthoFinder identified 20,547 orthogroups representing 97.9% of the genes across all 24 analyzed genomes (including the human outgroup). Of the orthogroups, 6,158 had all species present. In our *de novo* bank vole genome, there were 27,824 annotated genes, of which 84.2% were assigned to an orthogroup. Bank vole genes were present in 16,250 orthogroups. On average, the genomes included in the OrthoFinder analysis had 19,814 genes, with 98.2% of those assigned to orthogroups.

The completeness and large scaffold N50 (4.6 Megabases) of our bank vole assembly supported its inclusion in generating a Glires synteny network. Using the infomap clustering algorithm, we produced 19,694 microsynteny clusters from this overall synteny network. We did not expect dental genes to share the same microsynteny cluster, and instead examined whether each gene was in the same microsynteny cluster in species with rooted or unrooted molars. We identified 15 hierarchical orthogroups in which synteny was not conserved for at least half of the Glires with unrooted molars (Fig. 2). The genes form two groups (Fig. 2A), group 1, lacking synteny across Glires, and group 2, lacking synteny mainly in species with unrooted molars. Most of these genes also are missing from the orthogroups; only *Mmp20*, *Irx6*, *Aqp3*, *Sema3b*, and *Col4a1* were well represented in their orthogroups but not in their synteny networks (full comparisons of orthology and synteny are in Additional information 1). Overall, these genes represent multiple categories of “keystone” dental genes. Null mutations in keystone dental genes affect embryonic dental development (43): “shape” genes cause morphological errors; “eruption” genes prevent tooth eruption; “progression” genes stop the developmental sequence; “tissue” genes cause defects in tissues; “developmental process” genes are annotated with the “GO:0032502” gene ontology term; “dispensable” genes, while dynamically expressed in developing teeth, have no documented effect on phenotype; and “double” genes function redundantly with a paralog and only produce a phenotype when both genes are mutated. The group “other” is composed of the remaining protein coding genes (43). Most genes lacking conserved synteny in species with unrooted molars are in the “dispensable” category (Fig. 2D), thus the relationship between differences in these genes and tooth phenotypes is unclear, at least during embryonic development.

**Figure 2.**
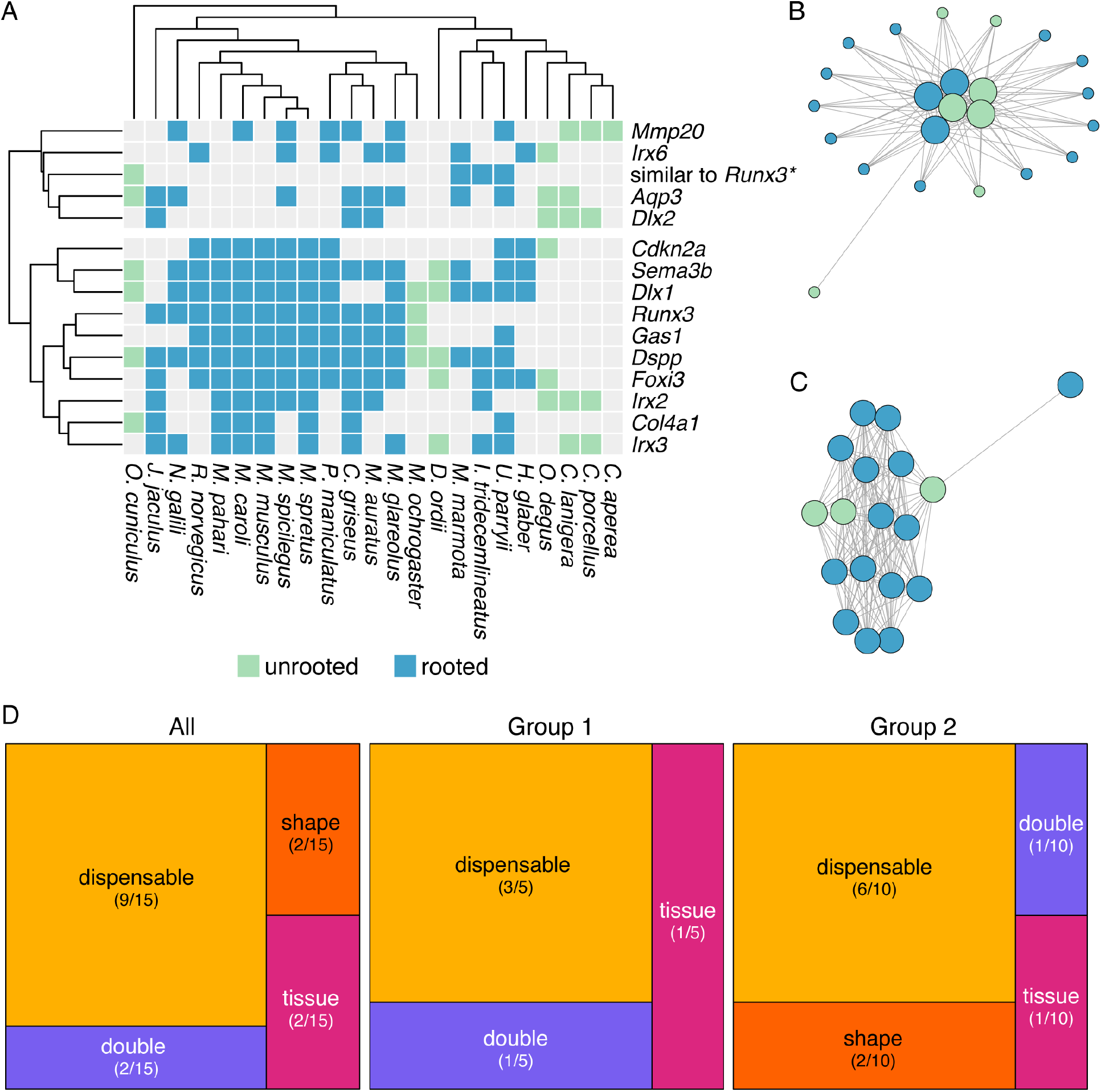
A. Presence (colored boxes) or absence (gray boxes) of gene sequences for each species in hierarchical orthogroups where fewer than half of the species with unrooted molars had conserved synteny. Columns are ordered according to phylogenetic positions (top) and rows are ordered by Euclidean distance clustering. Rows are split into two major groups: group 1, in which synteny is not conserved across Glires, and group 2, in which synteny is not conserved mainly in species with unrooted molars. * = One hierarchical orthogroup represented only four gene sequences annotated based on similarity to *Runx3*. **B** An example of a synteny network for genes in Group 1, displayed using the Fruchterman-Reingold layout algorithm in the R package *iGraph* (142). Small circles represent genes in the synteny network that are not part of the hierarchical orthogroup, large circles represent genes in the hierarchical orthogroup, and lines between circles represent a syntenic relationship between two species. Circle color represents whether species has rooted or unrooted molars following the same key in A. **C** An example synteny network for genes in Group 2, displayed using the Fruchterman-Reingold layout algorithm in the R package *iGraph* (142). Circles represent genes in the hierarchical orthogroup, and lines between circles represent a syntenic relationship between two species. Circle color represents whether species has rooted or unrooted molars following the same key in A. **D** Treemaps representing the keystone gene categories for all hierarchical orthogroups, the Group 1 hierarchical orthogroups, and the Group 2 hierarchical orthogroups. Most genes in each group are in the “dispensable” keystone gene category, which includes genes that are dynamically expressed during dental development but have no documented effect on phenotypes.

### Multiple dental genes under positive selection

We hypothesized that dental genes are undergoing positive selection in species with unrooted molars. Our positive selection analyses in PAML (phylogenetic analysis by maximum likelihood (44)) identified 6 dental gene orthogroups undergoing site-specific positive selection across Glires (Table 1). Four orthogroups with site-specific positive selection lacked synteny among Glires with unrooted molars: *Col4a1*, *Dspp*, *Runx3,* and the four-gene orthogroup with sequences similar to *Runx3* (Fig. 2A). We then assessed genes for site-specific positive selection in species with unrooted molars compared to species with rooted molars (branch-and-site-specific positive selection (45)), focusing on those genes with site-specific positive selection or evidence for loss of synteny. Two genes, *Dspp* and *Aqp1* were undergoing this branch-and-site specific positive selection. Both genes had a single highly supported site (posterior probability > 0.95) under positive selection in species with unrooted molars based on the Bayes Empirical Bayes method for identifying sites under selection implemented in PAML (46). *Dspp* also had multiple sites with moderate support (posterior probability > 0.75). The overall selection patterns on each gene differed. Maximum likelihood estimates of selection for *Dspp* showed the percentage of sites under purifying and neutral selection on all branches were nearly equal (47% and 44%, respectively). Percentages of sites under positive selection in the species with unrooted molars (foreground branches) were nearly evenly divided as well, with 5% of sites from branches where the species with rooted molars (background branches) were undergoing purifying selection and 4% of sites from branches where the species with rooted molars were under neutral selection. For *Aqp1*, nearly all sites were under purifying selection on all branches (91%), and few sites were under neutral selection on all branches (7%). Few sites were undergoing positive selection in the foreground branches and their distribution also was unevenly split between sites under purifying and neutral selection on background branches (0.6% and 0.04%, respectively). The complete list of dental genes with hierarchical orthogroups, microsynteny clusters, and positive selection test results are available in Additional file 1.

**Table 1.**
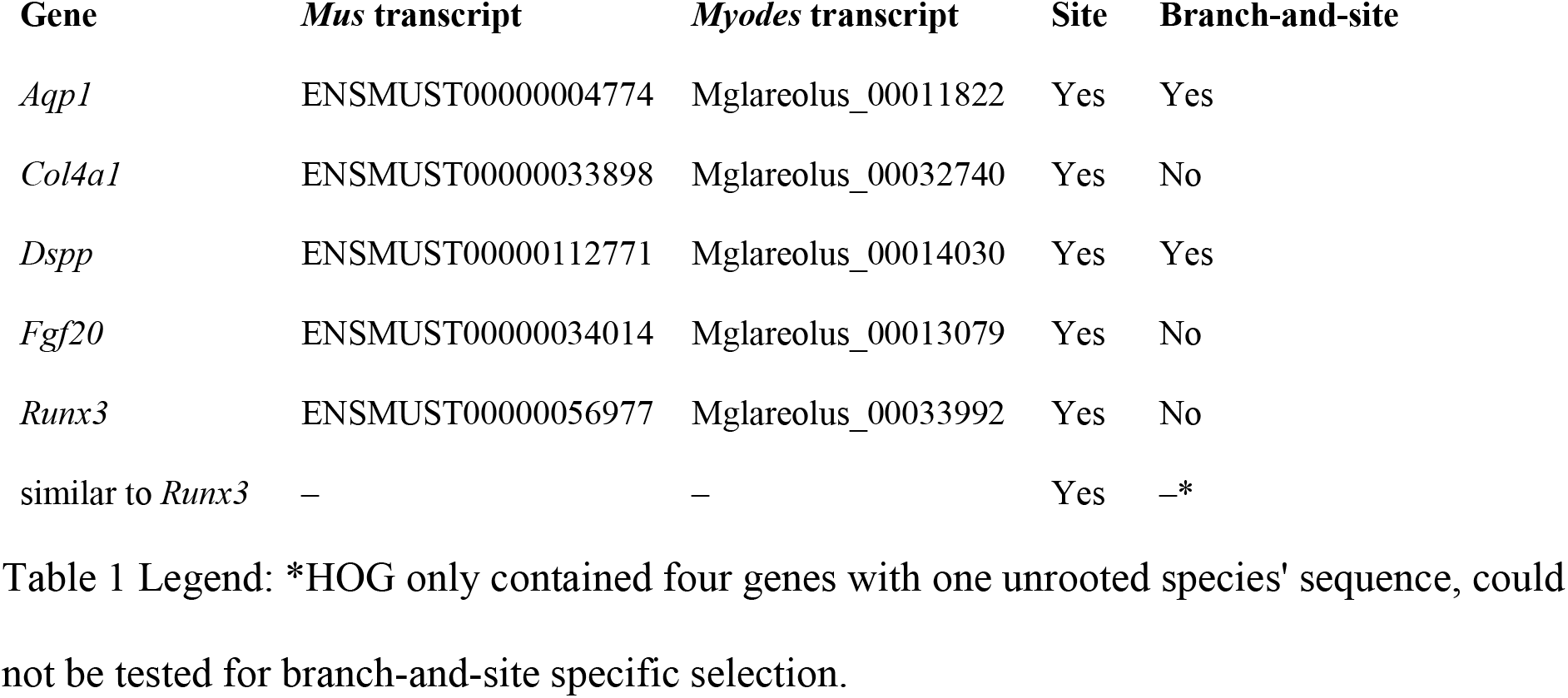
Genes undergoing site-specific and branch-and-site-specific positive selection.

Because genes under positive selection are often expressed at lower levels than genes under purifying selection (47–50), we also compared expression levels of *Dspp* and *Aqp1* in first molars (M1) at postnatal days 1, 15, and 21 (P1, P15, and P21) in bank voles (rooted molars) and prairie voles (unrooted molars) using quantitative PCR. Prairie vole molars expressed *Aqp1* at significantly lower levels than bank vole molars across all three ages (Fig. 3). Prairie vole P1 molars expressed significantly lower levels of *Dspp* than bank vole molars; at P15 and P21, their molars expressed *Dspp* at lower, but not statistically significantly different, levels than their bank vole equivalent. For both genes, the prairie vole had consistent expression levels across three biological replicates, while the bank vole had greater variation in expression levels across replicates.

**Figure 3.**
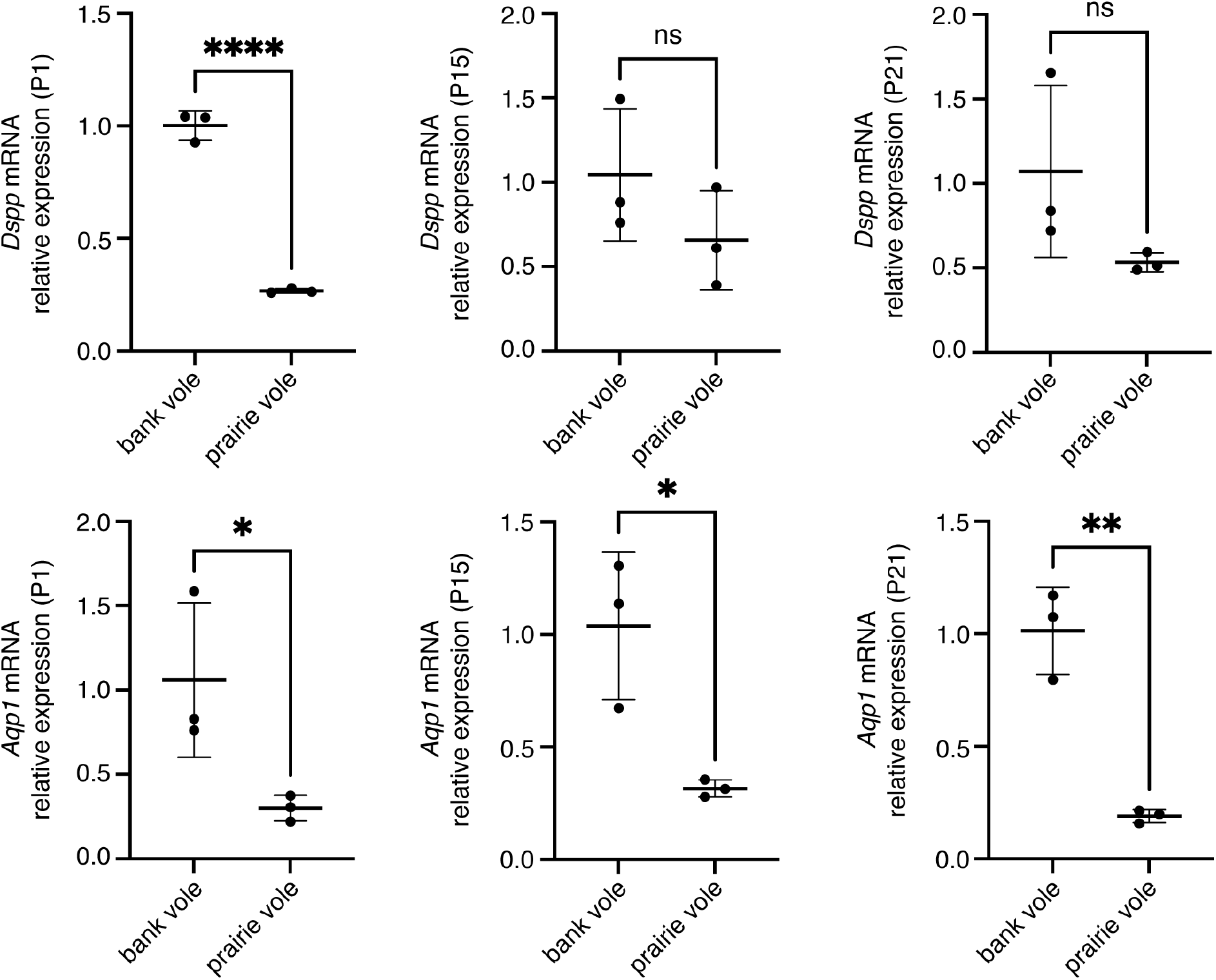
Quantitative PCR comparisons of *Dspp* and *Aqp1* expression between bank vole and prairie vole M1 at postnatal days 1, 15, and 21 (P1, P15, P21). Expression levels for both genes are lower in the prairie vole (unrooted molars), which supports the positive selection detected for these genes in species with unrooted molars.

### Few changes of secondary structure at positively selected sites

To detect whether substitutions at sites under positive selection influenced protein structure and evolution, we analyzed ancestral states and secondary structure across Glires. We first reconstructed ancestral sequences along the internal nodes of the Glires phylogeny for the genes undergoing branch-and-site specific positive selection to assess potential secondary structural changes in their protein sequences. At the best-supported site in *Dspp* (position 209 in the gapped alignment, Additional file 2), there were three major amino acid changes. The ancestral Glires sequence started with an asparagine (N) in this position. Two of the three species with unrooted molars represented in the *Dspp* dataset had amino acid substitutions at this position, with *Oryctolagus cuniculus* substituting a leucine (L) and *Dipodomys ordii* substituting an aspartic acid (D) at this position (Fig. 4A). All muroids (the clade including the voles in family Cricetidae and mice and rats in family Muridae) in our phylogeny substituted histidine (H) for the asparagine at this position. The secondary structure predicted at this position was a coil for most sequences but a helix for the *D. ordii* sequence (Fig. 5). *Aqp1* sequences varied greatly at the position under putative positive selection in species with unrooted molars (position 294 in the gapped alignment, Additional file 3). The ancestral state reconstruction showed twelve changes of the amino acid at this position across Glires (Fig. 4B), yet these changes did not affect the predicted secondary structure of the protein near this residue, which was a coil for all sequences tested. All secondary structure predictions are available in Additional file 4.

**Figure 4.**
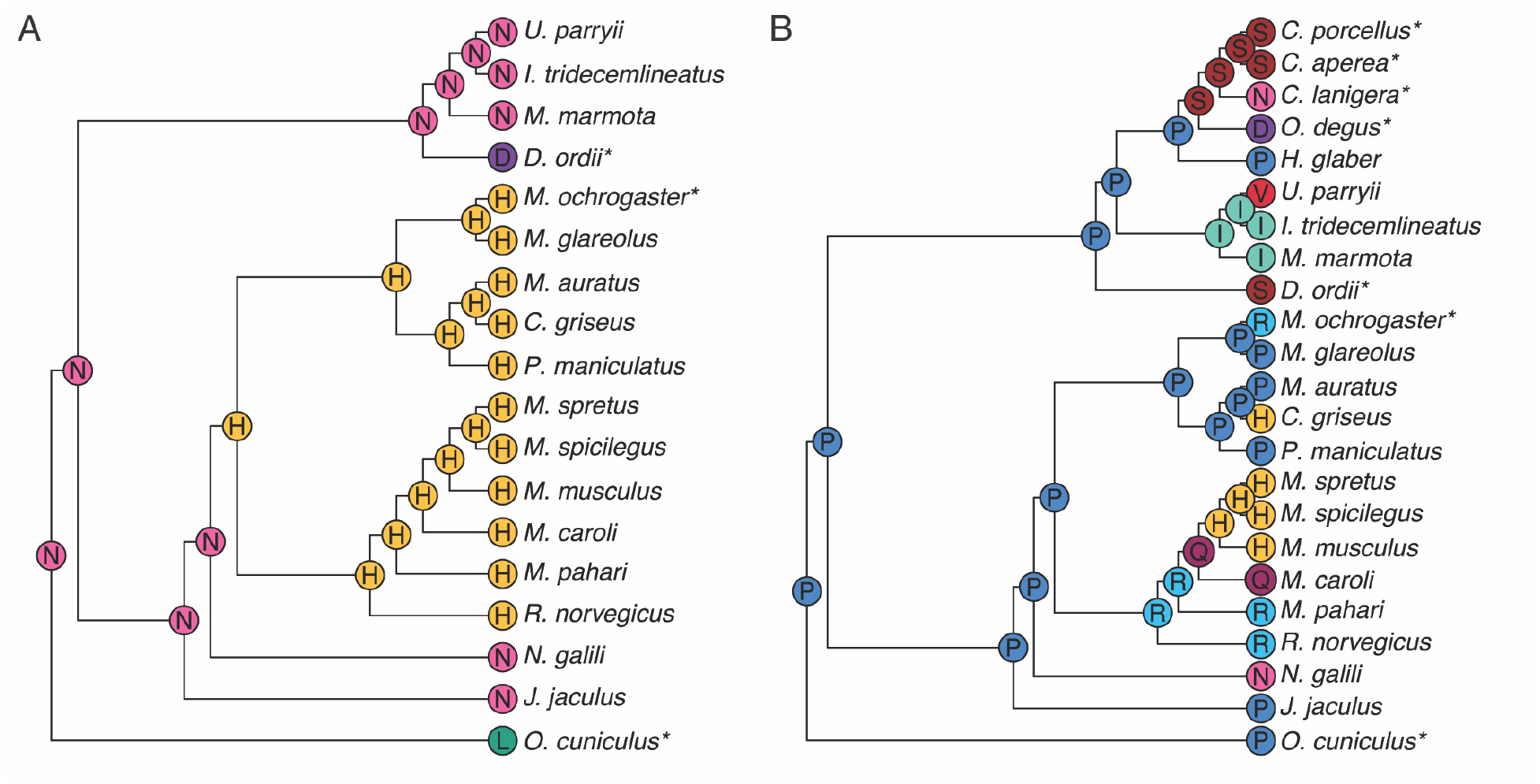
Ancestral state reconstructions of the residue under positive selection in PAML tests. Letters at tips and internal nodes represent IUPAC codes for amino acids and * denotes species with unrooted molars. **A** *Dspp*; **B** *Aqp1*.

**Figure 5.**
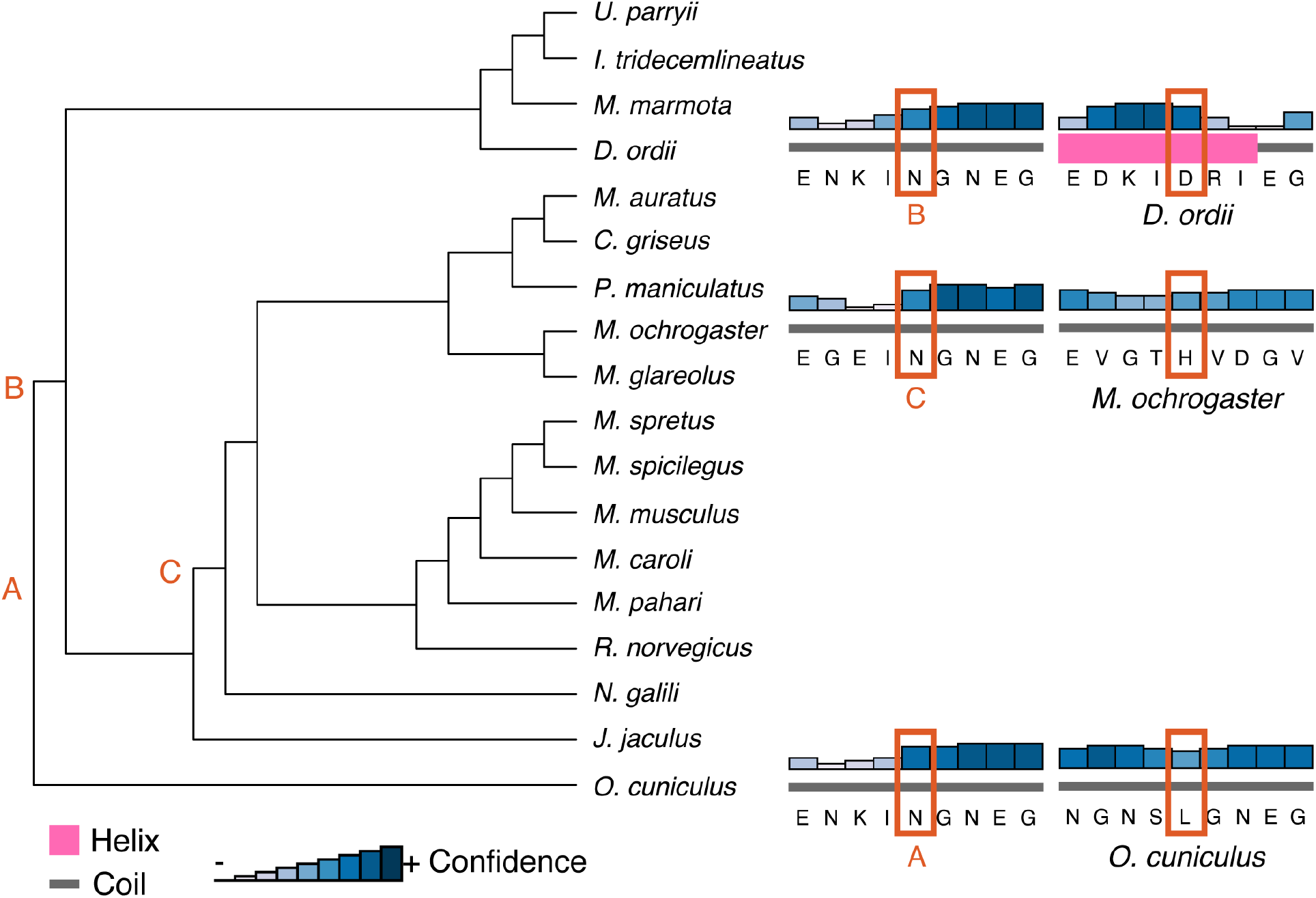
PSIPRED secondary structure predictions for the three species with unrooted molars represented in the *Dspp* sequences. Letters correspond to the most recent ancestor of each tip species where the amino acid at the site under positive selection differed: A, the predicted ancestor of *O. cuniculus*; B, the predicted ancestor of *D. ordii*; and C, the predicted ancestor of *M. ochrogaster*. Structure predictions, the relative confidence of the prediction, and the amino acid sequence for each pair of extant species and ancestor are on the right.

### Bank vole molar gene expression is similar to that of other Glires

We also assessed differential gene expression between mouse and bank vole molars across early development to study the effects of morphology on expression levels of dental genes. Our gene expression analysis focused on keystone dental gene categories. Our bank vole genome was like the mouse and rat genomes in terms of the numbers and expression patterns of genes annotated from these keystone categories (Table 2). Ordination of gene expression results from the bank vole and mouse data at embryonic day 13, 14, and 16 (E13, E14, E16) (43) by principal components analysis showed a distinct separation between the mouse and bank vole along the first principal component (PC1) of the 500 most variable genes (Fig. 6A). PC1 explained 82.81% of the variance in these genes; there are distinct, species-specific expression patterns in these tissues. Along PC2 (7.47% of variance explained), E13 and E14 samples differ from the E16 samples, although the difference in time points is much greater in bank voles. Ordination of just the keystone dental genes showed clear separations between tissues based on species and age (Fig. 6B). Within this focused set of genes, however, PC1 and PC2 explain less variance (44.8% and 28.84% respectively), and have a less clear relationship to species and age. There are two distinct, parallel trajectories for the mouse and bank vole. Although within each species there is separation by age along PC1 and PC2, mouse E16 and bank vole E13 occupy a similar position along PC1, and mouse E13 and bank vole E16 occupy a similar position along PC2.

**Figure 6.**
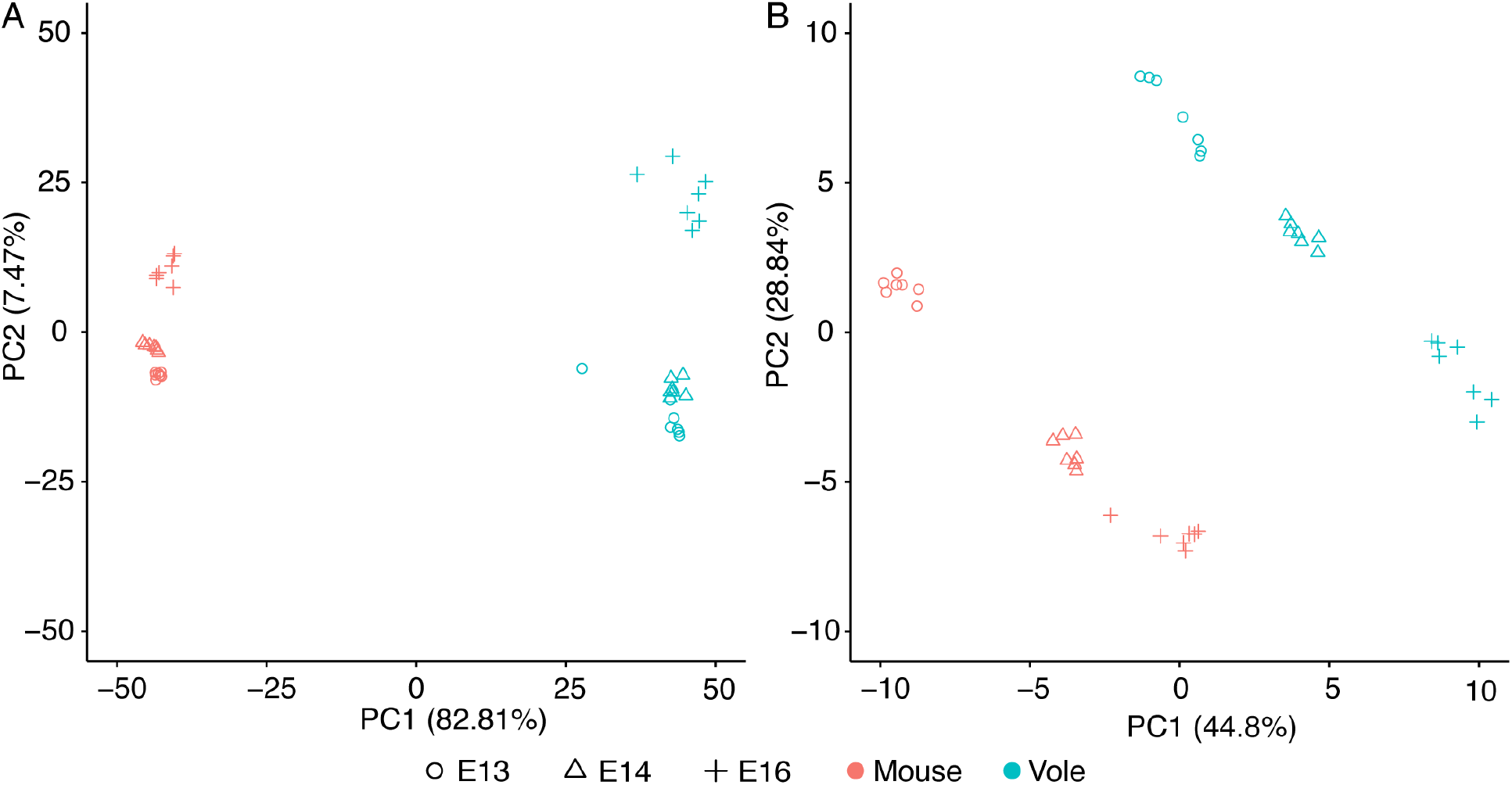
Principal component (PC) analyses of differentially expressed genes in mouse and bank vole M1. **A** PC1 and PC2 of the 500 most variable genes, showing a clear differentiation between species along PC1 and differentiation between age classes along PC2. **B** PC1 and PC2 of the keystone dental genes. Both PC1 and PC2 separate age classes within, but not between, the species, likely due to differences in developmental timing and molar morphology between mice and voles.

**Table 2.** P-values of permutation tests between keystone gene categories in bank vole M1 at embryonic days 13, 14, and 16.

Legend: Italicized values are statistically significant ( $p < 0.05$ )
|  | Tissue | Dispensable | Dev. Process | Other |
| --- | --- | --- | --- | --- |
| **E13 Progression** | *0.0310* | 0.0942 | *0.0436* | *0.0402* |
| **Shape** | 0.6431 | 0.9041 | 0.2289 | 0.0995 |
| **Double** | 0.1292 | 0.1521 | 0.0716 | 0.0655 |
| **E14 Progression** | *0.0136* | *0.0383* | *0.0437* | *0.0401* |
| **Shape** | 0.3115 | 0.4725 | 0.0922 | *0.0454* |
| **Double** | 0.1288 | 0.0945 | 0.0709 | 0.0630 |
| **E16 Progression** | *0.0140* | *0.0401* | *0.0303* | *0.0274* |
| **Shape** | 0.3770 | 1 | 0.1831 | 0.0662 |
| **Double** | 0.1343 | 0.1099 | 0.0638 | 0.0596 |

Examining individual genes underlying the differences between mouse and vole molars, we note several upregulated genes in our vole molars are broadly expressed in developing molars of other vole species (51,52). Relative to the mouse molars, vole molars overexpressed genes related to forming tooth cusps, including *Bmp2*, *Shh*, *p21* (also known as *Cdkn1a*), and *Msx2*, a difference explained by the faster patterning and larger number of cusps in the vole molar compared to the mouse molar (52). Another gene upregulated in the patterning stage vole molar is *Fgf10*, which is associated with delayed root formation later in vole molar development (9).

Nevertheless, developing bank vole molars at E13, E14, and E16 expressed keystone dental genes in overall proportions like those observed at analogous stages of mouse and rat molar development (Fig. 7). Permutation tests within each bank vole sample showed that log counts for the set of genes related to the progression of dental development were significantly higher than those in the tissue, dispensable, developmental process, and “other” categories at E14 and E16. The progression gene counts in E13 molars were higher for all of these except the dispensable category. Shape category genes also were significantly higher than “other” category genes in the E14 tissue. Overall, even though we observed conserved expression patterns of dental genes at the system level, individual genes involved in cusp patterning and morphology differed between the mouse and the vole.

**Figure 7.**
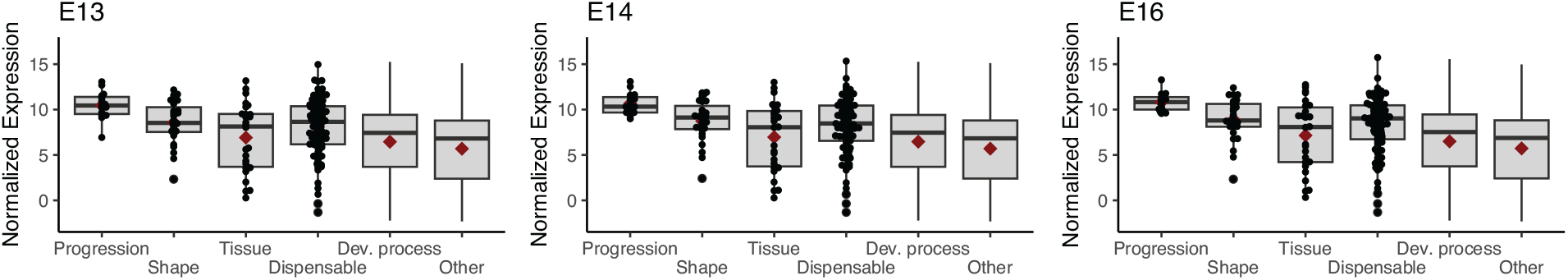
Box and whisker plots showing normalized log base 2 expression levels for each keystone gene category in bank vole M1 at embryonic days 13, 14, and 16. Horizontal bar and diamond within each box represent the median and mean values. Individual datapoints are displayed for smaller keystone gene categories. Gene expression profiles at these stages are comparable to mouse and rat molars at analogous developmental stages, as seen in Hallikas et al. 2021.

## DISCUSSION

Our two goals in sequencing the genome of *Myodes glareolus* were to support the development of a comparative system for studying tooth root development and to investigate the evolution of dental genes in Glires, a clade in which ever-growing molars have evolved multiple times (1). Our new *M. glareolus* assembly and annotation captured nearly all of the single-copy orthologs for Euarchontoglires and provided scaffolds with sufficient length for synteny analyses. It was well represented in ortholog groups and microsynteny clusters across Glires. We tested the hypothesis that dental genes are undergoing site-specific positive selection in species with unrooted molars (branch-and-site specific positive selection (45)). We predicted that lack of conserved syntenic relationships in species with unrooted molars could place dental genes in regulatory and selective environments that promote changes among genes relevant to tooth root formation. Our analyses identified 15 dental genes without conserved syntenic relationships across Glires and two dental genes, *Dspp* and *Aqp1*, under positive selection in species with unrooted molars. We also demonstrated conserved patterns of gene expression among dental keystone genes between bank voles and mice during early embryonic development, and deviations from these conserved patterns likely related to differences in molar morphology between the two species.

We identified 15 genes which were not syntenic in at least half of the species with unrooted molars, and six genes undergoing site-specific positive selection across all Glires. Although four of the orthogroups with site-specific positive selection lacked synteny in species with unrooted molars, only *Col4a1* was well represented among these species in its orthogroup. The two genes undergoing branch-and-site-specific positive selection in species with unrooted molars, *Dspp* and *Aqp1*, both maintained their synteny relationships across the Glires studied. Although we predicted loss of synteny for dental genes in Glires with unrooted molars could result in sequence evolution by placing genes in new selective contexts, our analyses did not support a strong relationship between non-syntenic genes and branch-and-site-specific positive selection. Maximum likelihood estimates of selection on each site for the genes with branch-specific positive selection revealed different overall selective pressures on *Dspp* and *Aqp1*; *Dspp* sites on background branches (i.e., branches with species that have rooted molars) were under a mix of purifying and neutral selection, while nearly all *Aqp1* background branch sites were under purifying selection. These selection regimes suggest there is greater conservation for *Aqp1* function across Glires than for *Dspp* function. Gene duplication can result in functional redundancy and evolution toward a novel function in some genes (53–56), which may explain positive selection in *Aqp1*, as there are other aquaporin family genes present. Although *Dspp* has no paralogs, it overlaps functionally with other SIBLING family proteins (e.g., *Opn*, *Dmp1*) (57,58).

*Aqp1* and *Dspp* play different functional roles during dental development. Under the keystone dental development gene framework, *Aqp1* is a “dispensable” gene: developing teeth express it, but tooth phenotypes do not change in its absence. *Aqp1* is expressed in endothelia of microvessels in the developing tooth (59,60). *Dspp* may be particularly relevant for the formation of an unrooted phenotype if its expression domain or function have been modified in species with unrooted molars. *Dspp* is a “tissue” category keystone dental gene, meaning the main effects of a null mutation occur during the tissue differentiation stage of dental development (43). Null mutations of *Dspp* cause dentin defects in a condition called dentinogenesis imperfecta (61,62); in some patients, teeth form short, brittle roots (62,63). *Dspp* knockout mice also exhibit the shortened root phenotype, among a variety of other defects in both endochondral and intramembranous bone, due to the disruption of collagen and bone mineralization (64–66).

Our ancestral sequence reconstructions and estimated secondary protein structures allowed us to assess whether nonsynonymous substitutions at sites under positive selection resulted in structural differences, thus potentially affecting protein function. Although unrooted molars are a convergent phenotype across Glires, the sites under positive selection did not converge on the same amino acid substitution in species with unrooted molars, and *Aqp1* appeared particularly labile at this residue. The non-synonymous substitutions at these sites often resulted in changes of properties of the amino acid in the sequence, for example in *Dspp*, polar asparagine was replaced with non-polar leucine in *O. cuniculus*. Only one of these substitutions changed the predicted secondary structure. Nevertheless, single amino acid substitutions do produce dental phenotypes for both *Dspp* (67) and *Aqp1* (68), thus we cannot rule out functional changes in these genes in species with unrooted molars.

Although the exact relationship between gene expression and sequence divergence remains unclear (69), studies of genome evolution across small numbers of mammal species show correlations between gene sequence divergence and levels of expression (70). In particular, highly-expressed genes are more likely to experience purifying selection (47–50), while lowly-expressed genes and tissue-specific genes may experience positive selection (48). The decreased expression of *Dspp* and *Aqp1* in prairie vole M1 compared to that of the bank vole M1 thus supports our finding of positive selection in these genes in species with unrooted molars. If all species with unrooted molars also exhibit decreased expression levels of *Dspp* and *Aqp1*, it could suggest a strong link between lower levels of the genes and the unrooted phenotype.

Without analyses of functional variation caused by positive selection at these coding sites, or spatial sampling to determine where these genes may be expressed during development, we are limited from exploring the specific effects of *Dspp* and *Aqp1* on root formation. Nevertheless, we found evidence for evolution of these genes in Glires with unrooted molars, and *Dspp* especially has clinical relevance for tooth root formation. Future studies should explore the spatial distribution of *Dspp* expression, which could be relevant to functional changes in Glires with unrooted molars. If positive selection and corresponding amino acid changes identified in *Dspp* here modify its expression domain or its interaction with yet-unidentified root formation co-factors, it may serially reproduce the unrooted incisor phenotype in molars.

Our RNA sequencing results supported the bank vole as a suitable system for studying dental development. Although molar morphology differs considerably across mammals, candidate-gene approaches have identified numerous conserved genes involved in tooth development and morphological patterning (71). Studies of single genes or gene families have identified shape-specifying roles common to multiple species (52,72–74), and high-throughput sequencing of mouse and rat molars demonstrate that both species express sets of dental development genes in similar proportions during early stages of tooth development (43). The similarity of our high-throughput RNA sequencing results (Fig. 7) to the mouse and rat results in previous studies suggest overall expression patterns of keystone dental development genes within each stage are conserved across Glires. Our principal component analyses and differential expression analyses measuring changes between mouse and bank vole molars, however, showed that several dental genes’ expression levels differed significantly by species and age. Previous research has documented organ expression patterns that are conserved across species early in development and diverge over time, with some major organs displaying heterochronic shifts in some species (75). If the major source of variation in keystone dental gene expression patterns between mice and bank vole molars were solely attributable to species, we might expect to see clear separation between the species along the first or second principal component (PC1 or PC2), like that observed in PC1 of the 500 most variable genes (Fig. 6). If molar development follows the diverging expression patterns observed in other organs, we might expect just the earliest age classes to align on one, or multiple, PCs. Instead, we found two trajectories that were nearly parallel across PC1 and PC2 and multiple keystone dental genes that were significantly differentially expressed with respect to species and age. This variation between species is likely driven by the larger number of cusps in the vole molar, and corresponding upregulation of genes regulating cusp formation. The overall acceleration of patterning in vole molars likely explains the significance of the age variable in our expression results, causing a heterochronic shift in the expression patterns.

Our analyses were limited by the small number of rodent species with sufficiently annotated genomes to be included in synteny and positive selection analyses. This limitation left us with a small phylogeny for our ancestral state reconstructions, which thus did not encompass the full diversity of Glires tooth roots, and potentially weakened model-based genomic analyses. Although positive selection analyses using the Bayes Empirical Bayes criterion are robust to smaller sample sizes (46), incomplete sampling can affect estimations of ancestral characteristics (76). Innovations in paleoproteomics also offer the opportunity to compare fossil species’ dental gene sequences directly to living and estimated ancestral sequences (77,78). By incorporating data for extinct Glires in both morphological and molecular analyses, we can further elucidate links between dental gene evolution and unrooted teeth.

## CONCLUSIONS

Our genomics and transcriptomics analyses, based on our newly sequenced, high-quality draft bank vole genome assembly and annotation, showed that bank vole early tooth development is comparable to other commonly used rodent models in dental development research. We identified 6 dental gene orthogroups that were undergoing site-specific positive selection across Glires and two genes, *Dspp* and *Aqp1*, that were undergoing site-specific positive selection in Glires with unrooted molars. *Dspp* appears particularly relevant to root formation, as loss-of-function mutations cause a dentin production defect that can result in shortened tooth roots. Future research must explore the functional role that *Dspp* plays in tooth root formation in Glires and other clades. The rodent dentary is an exciting system for understanding tooth development; it provides an easily manipulated set of tissues that can be produced quickly and features a lifelong population of stem cells in the incisor with genomic mechanisms that are potentially replicated across other teeth in species with unrooted molars. Our results identify candidate genes for future analyses, and our draft bank vole genome and annotation improve the utility of this species for comparative dental research that can uncover the genetic mechanisms of tooth root formation.

## METHODS

### Tissue collection and sequencing

To assemble the bank vole genome, we sequenced tissues from a single adult male specimen housed in a colony at the UCSF Mission Center Animal Facility. We euthanized the animal according to UCSF IACUC protocol AN189916 and harvested muscle, kidney, heart, and liver tissue, which were immediately frozen at −80°C. Tissues were sent to a third-party sequencing service, where they were combined and homogenized to achieve appropriate mass for high molecular weight DNA extraction. We targeted 60x coverage with 150 base pair (bp) reads using 10X Chromium linked-read chemistry (79,80) sequenced on the Illumina platform. We also targeted 10x coverage with Pacific Biosciences SMRT long-read chemistry. For genome annotation and gene expression analyses, we collected seven biological replicates each of first molars at embryonic days 13-16 (E13, E14, E15, E16), second molars at E16, and jaw tissues at E14 under University of Helsinki protocols KEK16-021, KEK19-019, and KEK17-030 and stored them in RNAlater at −80°C for RNA sequencing, following a tissue harvesting protocol established for mice and rats (43). We extracted RNA from these tissues using a guanidium thiocyanate and phenol-chloroform protocol combined with an RNeasy column purification kit (Qiagen) based on the keystone dental gene protocol (43). Single-end 84 bp RNA sequencing was performed using the Illumina NextSeq 500 platform.

### Genome assembly and quality control

We first assembled only the 10X Chromium linked reads using the default settings in Supernova 2.1.1. (79,80). We selected the “pseudohaplotype” (pseudohap) output format, which randomly selects between potential alleles when there are two possible contigs assembled for the same region. This option produces two assemblies, each with a single resolved length of the genome sequence (79–81). We used our lower-coverage, long-read data for gap filling and additional scaffolding. First, we estimated the genome’s length using the raw sequence data in GenomeScope (82), which predicted a length of 2.6 gigabases. We then performed error correction of the long reads using Canu (83), removing reads shorter than 500 base pairs (bp) and disregarding overlaps between reads shorter than 350 bp. We kept only those reads with minimum coverage of 3x for scaffolding. Following long read error correction, we used Cobbler and RAILS (84) with a minimum alignment length of 200 bases to accept matches for gap filling and scaffolding of both pseudohap assemblies.

For quality control, we assessed both unscaffolded and long-read scaffolded pseudohap assemblies by standard assembly length statistics with QUAST (85) and presence of single-copy orthologs with BUSCO v3 (86). Both scaffolded assemblies were approximately 2.44 Gigabases long, with an N50 (the length of the shortest scaffold at 50% of the total assembly length) of 4.6 Megabases; we refer to them as Pseudohap1+LR and Pseudohap2+LR. The Pseudohap1+LR assembly had 17,528 scaffolds over 1000 bp long, and the Pseudohap2+LR assembly had 17,518 scaffolds over 1000 bp long (Table 3). BUSCO searched for universal single-copy orthologs shared by Euarchontoglires, recovering 89.4% of these genes in the scaffolded Pseudohap1+LR assembly and 92.8% of the single-copy orthologs in the scaffolded Pseudohap2+LR assembly (Fig. 8). The two assemblies were similar length and contiguity, but we based annotation and downstream analyses on Pseudohap2+LR because it recovered more single-copy orthologs.

**Figure 8.**
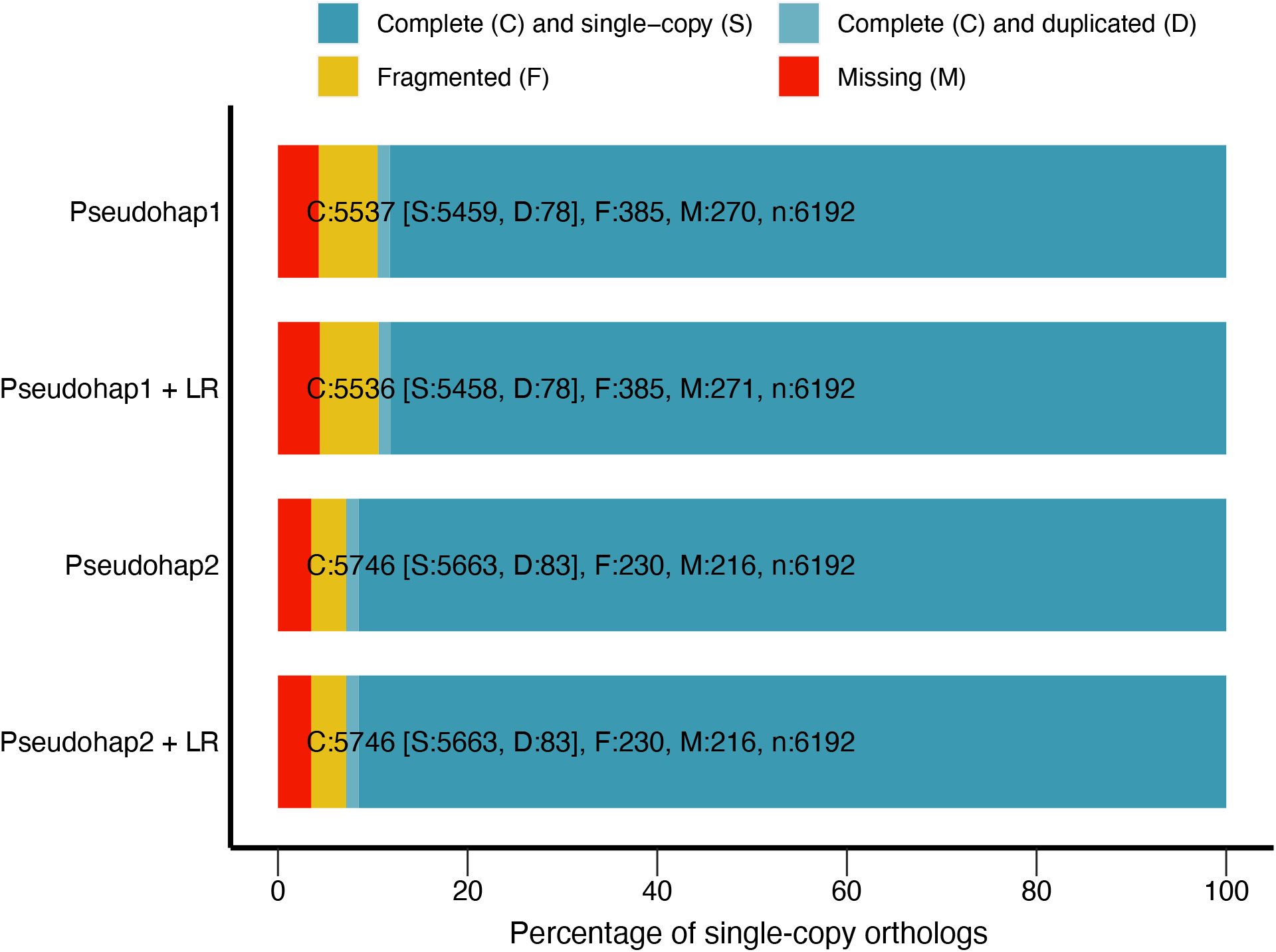
BUSCO single-copy ortholog recovery for each “pseudohaploid” version of our draft bank vole genome assembly and these version after long-read scaffolding (denoted by “+ LR”). Each bar represents the cumulative proportion of the 6,192 single-copy orthologs for Euarchontoglires identified by BUSCO represented by complete single-copy, complete-duplicated, fragmented, and missing orthologs. The Pseudohap2 and Pseudohap2 + LR assemblies had the best single-copy ortholog recovery.

**Table 3.** QUAST assembly statistics for *de novo* bank vole (*Myodes glareolus*) genome.

Legend: \*assembly used for annotation and downstream analyses in this paper.
|  | Pseudohap1 | Pseudohap1+LR | Pseudohap2 | Pseudohap2+LR* |
| --- | --- | --- | --- | --- |
| **Largest contig** | 27939478 | 32658832 | 27937749 | 32657565 |
| **Total length** | 2434151515 | 2441426554 | 2434099357 | 2441472313 |
| **GC (%)** | 41.88 | 41.89 | 41.88 | 41.89 |
| **N50** | 4187179 | 4579815 | 4187179 | 4558134 |
| **N75** | 1689669 | 1818134 | 1687188 | 1810460 |
| **L50** | 170 | 153 | 170 | 154 |
| **L75** | 388 | 357 | 388 | 358 |
| **Ns per 100 kbp** | 1151.99 | 1030.75 | 1151.96 | 1030.48 |

### Genome annotation

We annotated the genome using multiple lines of evidence in three rounds of the MAKER pipeline (87–89). For evidence from gene transcripts, we assembled a *de novo* transcriptome assembly of the single-end RNA sequences pooled from all molar and jaw tissues using Trinity (90). We also included cDNA sequences from the *Mus musculus* assembly GRCm38 to provide additional transcript evidence from a close relative with a deeply annotated genome. We used SwissProt’s curated protein database to identify protein homology in the genome. Two libraries of repeats provided information for repeat masking: the Dfam Rodentia repeat library (91–93) and a custom library specific to the bank vole estimated with a protocol modified from Campbell et al. (88). The custom library features miniature inverted-repeat transposable elements identified with default settings in MiteFinder (94), long terminal repeat retrotransposons extracted with the GenomeTools LTRharvest and LTRdigest functions (95) based on the eukaryotic genomic tRNA database, and *de novo* repeats identified with RepeatModeler (96). We combined elements identified by these programs into a single repeat library, then removed any elements that matched to a custom SwissProt curated protein database excluding known transposons. The custom repeat library is available in Additional file 5. We trained a custom gene prediction model for MAKER as well. The first iteration of the model came from BUSCO’s implementation of augustus (97). Between each round of MAKER annotation, we further updated the gene prediction model with augustus.

MAKER considered only contigs between 10,000-300,000 bp long during annotation. Our second and third iterations of MAKER used the same settings but excluded the “Est2genome” and “protein2genome” functions, as recommended in the MAKER tutorial. We included a SNAP (98) gene prediction model based on the output of the first round of annotation during the second and third iterations of MAKER annotation. Annotation quality (i.e., agreement between different lines of evidence and the MAKER annotation) was assessed visually in JBrowse after each iteration and using *compare_annotations_3.2.pl* (99), which calculates the number of coding and non-coding sequences in the annotation in addition to basic statistics about sequence lengths. Our MAKER annotation covered 2.41 Gb of the scaffolded Pseudohap2 assembly in 4,125 scaffolds. These scaffolds contained 27,824 coding genes (mRNA) and 15,320 non-coding RNA sequences. The average gene length was 12,705 bp. Most annotations (91.4%) had an annotation edit distance (AED) of 0.5 or better. AED is a measure of congruency between the different types of evidence for an annotation, where scores closer to zero represent better-annotated genes (100).

### Orthology and synteny analyses

We analyzed orthology and synteny of the bank vole genome to understand gene and genome evolution related to dental development across Glires with rooted and unrooted molars. We obtained genomes from Ensembl for 23 Glires species and one phylogenetic outgoup, *Homo sapiens* (Table 4). These genomes all had an N50 over 1 Mb, which improves synteny assessment (101). We first analyzed all 24 genomes for groups of orthologous genes (orthogroups) in OrthoFinder (102), providing a tree topology based on the Ensembl Compara reference tree (Fig. 1) to guide orthology detection. Because we would not analyze the human outgroup in downstream analyses, we implemented the OrthoFinder option that splits orthogroups at the root of Glires (hierarchical orthogroups), thus any group of orthologs studied here represents only genes with shared, orthologous evolutionary history within Glires. We selected MAFFT (103) for multiple sequence alignment and fastme (104) for phylogenetic tree searches within OrthoFinder. We retained the gene trees estimated for each orthogroup for downstream analyses.

**Table 4.** Genomes used in orthology, synteny, and positive selection analyses.

Legend: \*Species with unrooted molars; \*\*Peptide annotation used as outgroup only in
| Species | Assembly | Citation |
| --- | --- | --- |
| *Myodes glareolus* | CUNY_Mgla_1.0 | This paper |
| *Cavia porcellus* * | Cavpor3.0 | (122) |
| *Cavia aperea* * | CavAp1.0 | (123) |
| *Marmota marmota* | marMar2.1 | (124) |
| *Microtus ochrogaster* * | MicOch1.0 | (125) |
| *Mus musculus* | GRCm39 | (126) |
| *Oryctolagus cuniculus* * | OryCun2.0 | (122) |
| *Dipodomys ordii* * | Dord_2.0 | (122) |
| *Jaculus jaculus* | JacJac1.0 | (127) |
| *Rattus norvegicus* | Rnor_6.0 | (128) |
| *Mus pahari* | PAHARI_EIJ_v1.1 | (129) |
| *Mus caroli* | CAROLI_EIJ_v1.1 | (129) |
| *Mus spretus* | SPRET_EiJ_v1 | (130) |
| *Mus spicilegus* | MUSP714 | (131) |
| *Cricetulus griseus* | CHOK1GS | (132) |
| *Mesocricetus auratus* | MesAur1.0 | (133) |
| *Peromyscus maniculatus* | HU_Pman_2.1 | (134) |
| *Nannospalax galili* | S.galili_v1.0 | (135) |
| *Octodon degus* * | OctDeg1.0 | (136) |
| *Heterocephalus glaber* (F) | HetGla_female_1.0 | (137) |
| *Chinchilla lanigera* * | ChiLan1.0 | (138) |
| *Urocitellus parryi* | ASM342692v1 | (139) |
| *Ictidomys tridecemlineatus* | SpeTri2.0 | (140) |
| *Homo sapiens* ** | GRCh38 | (141) |

OrthoFinder analysis.

Although dental development genes are spread throughout the genome, we were interested in whether each gene remained in the same local arrangement across species of Glires. We prepared each genome annotation and sequence file for synteny analysis using the reformatting functions of Synima (105) to extract each peptide sequence associated with a gene coding sequence in the Ensembl annotation. Collinear synteny blocks estimated by MCScanX (106) formed the basis for synteny network analyses using the SynNet pipeline (107–109). We inferred networks from the top five hits for each gene, requiring any network to have a minimum of 5 collinear genes and no more than 15 genes between a collinear block, settings that perform well for analyzing mammal genomes (109). Using the infomap algorithm, we clustered the synteny blocks into microsynteny networks, from which we extracted network clusters corresponding to the list of keystone dental genes (43). For each dental gene hierarchical orthogroup, we assessed whether genes of species with unrooted molars were missing from the synteny networks that contained other Glires species’ sequences, representing loss of synteny for those species.

### Positive selection analysis

We aligned protein sequences for each dental gene orthogroup with clustal omega (110) using default settings. Based on universal translation tables, we obtained codon-based nucleotide alignments with pal2nal (111), removing sites in which any species had an indel (i.e., ungapped) and formatting the output for analysis in PAML (44). We pruned and unrooted the orthogroup gene trees from OrthoFinder to contain only tips representing the genes in each synteny network or orthogroup under analysis in PAML. We tested whether any of the genes were undergoing positive selection using a likelihood ratio test comparing site-specific models of “nearly neutral” and positive selection. In these models, μ, the ratio of nonsynonymous to synonymous nucleotide substitutions (also known as dN/dS), can vary at each codon site. In the “nearly neutral” model, μ can take values between 0 and 1, while the positive selection model allows sites to assume μ values greater than 1 (46,112). We estimated κ (the ratio of transitions to transversions) and μ from initial values of 1 and 0.5, respectively, for both tests.

Dental genes with significant site-specific positive selection or those lacking synteny in species with unrooted molars formed the basis for our second set of positive selection tests using a branch-and-site model of positive selection. This model allows μ to vary not only among codon sites, but also between “foreground” and “background” lineages (46). We marked the species with unrooted molars as foreground lineages, then ran the model twice: once with μ unconstrained to detect sites undergoing positive selection only on foreground branches, and a second time and with μ fixed to 1, or neutral selection. A likelihood ratio test of the two models determined whether the lineage-specific positive selection model was more likely than a neutral model, and Bayes Empirical Bayes analyses (46) produced posterior probabilities to identify sites under positive selection.

Genes under positive selection also tend to have lower expression levels (48), thus we compared expression of the genes with branch-and-site specific positive selection between the prairie (unrooted molars) and the bank vole (rooted molars) to provide further support for selective differences. We collected three biological replicates of first molars from both species at three postnatal stages (P1, P15, and P21) and immediately preserved them at −80°C in lysis buffer (Buffer RLT; Qiagen) supplemented with 40 μM dithiothreitol. RNA was extracted from homogenized tissues using a RNeasy column purification kit (Qiagen). We assessed concentration and purity of extracted RNA using a NanoDrop 2000 spectrophotometer (ThermoFisher Scientific). Using 1 μg of RNA, we synthesized cDNA using a high-capacity cDNA reverse transcription kit (ThermoFisher Scientific). We used 1 μL diluted cDNA (1:3 in ddH_2_O) and iTaq Universal SYBR Green Supermix (Bio-rad) in the Bio-rad CFX96 real-time PCR detection system for qPCR experiments, producing three technical replicates for each biological replicate. We normalized cycle threshold (CT) values of genes of interest to GAPDH expression levels and calculated relative expression levels as 2^-ΔΔCT^. A two-tailed unpaired t-test calculated in Prism 9 measured whether expression of these genes significantly differed between bank voles and prairie voles. The oligonucleotide primers for each species and gene are in Additional file 6.

### Sequence and secondary structure evolution

We performed ancestral sequence reconstruction on the codon sequences of the genes that had evidence of branch-and-site specific positive selection to understand how the sequence has changed through time. The gapped clustal omega alignments were the basis for ancestral sequence reconstruction on the Glires species tree (Fig. 1) using pagan2 (113). For each gene, we plotted amino acid substitutions at the site with potential positive selection. Finally, we predicted secondary structures (i.e., helices, beta sheets, and coils) for each unrooted species’ protein sequence and the reconstructed ancestral sequence prior to the change at the site under positive selection using the PSIPRED 4.0 protein analysis workbench (114,115). Comparing these predictions across the phylogeny, we assessed how these substitutions at the site under selection may affect the structure of each protein.

### Developmental gene expression

We performed quality control and filtering of the short reads for the seven replicates of first molar tissues at E13, E14, and E16 using the nf-core/rnaseq v. 3.11.2 workflow (116) for comparability to previous mouse and rat analyses (43). RNAseq reads were evaluated and adapter sequences were filtered using FastQC v. 0.11.9 (117) and Cutadapt v. 3.4 (118), and ribosomal RNA was removed using SortMeRNA v. 4.3.4 (119). We then aligned trimmed sequences to our bank vole annotation using Salmon v. 1.10.1 (120). Counts were then normalized by gene length. We categorized gene count data into functional groups based on their established roles in tooth bud development (43) using the one-to-one orthology list between our bank vole genome and the mouse GRCm39.103 genome annotation generated from our OrthoFinder output. Using the rlog function of DESeq2 (121), we normalized gene counts within each functional group on a log2 scale. A permutation test assessed whether the mean counts of the progression, shape, and double functional groups were significantly different from genes in the tissue, dispensable, and “other” groups (which are potentially relevant later in development) based on 10,000 resampling replicates of the dataset (43).

We also assessed differential expression between the bank vole first molar and published mouse M1 data at the same three time points (GEO accession GSE142199 (43)), combining the data based on the one-to-one orthology relationships used in the functional permutation analysis. Using the mouse E13 molar as the reference level, we modeled expression as a response to species (mouse or vole), embryonic day (E13, E14, or E16), and the interaction between species and day. We considered as significant any gene with a log fold change greater than 1, log fold change standard error less than 0.5, and false discovery rate adjusted p value less than 0.05.

## Supporting information

Additional file 1: Dental gene results

Additional file 2: Dspp gapped alignment

Additional file 3: Aqp1 gapped alignment

Additional file 4: Structure predictions

Additional file 5: Custom repeat library

Additional file 6: Oligonucleotide primers

## DECLARATIONS

## Ethics approval

The University of California, San Francisco (UCSF) Institutional Animal Care and Use Program and the Finnish national animal experimentation board approved protocols for humane euthanasia and collection of tissues for animals used in this study under protocols AN189916 (UCSF) and KEK16-021, KEK19-019, and KEK17-030 (University of Helsinki).

## Availability of data and materials

The datasets supporting the conclusions of this article are available in the GenBank repository under the BioProject PRJNA1050237 (genome accession number JBBHLL000000000) and in the article’s additional files.

## Competing interests

The authors declare that they have no competing interests.

## Funding

This research was supported by National Science Foundation grants CNS-0958379, CNS-0855217, OAC-1126113, and OAC-2215760 through the City University of New York High Performance Computing Center at the College of Staten Island; OAC-1925590 through the MENDEL high performance computing cluster at the American Museum of Natural History; Academy of Finland to JJ; Doctoral Programme in Biomedicine, University of Helsinki to MMC; and National Institutes of Health NIDCR R01-DE027620 and R35-DE026602 to ODK.

## Authors’ contributions

ZTC and ODK designed the study. ZTC and PM performed animal husbandry. ZTC performed and oversaw tissue sampling, sequencing, genome assembly and annotation for *Myodes glareolus*. ZTC, AS, EC, and MA performed genome computational analyses. PL performed qPCR analyses. OH, MMC, RDR, and JJ designed and implemented RNA sequencing experiments. ZTC wrote and all authors contributed to and approved the manuscript.

## Acknowledgements

The authors thank A. Joo, N. Ahituv, G. Amato, A. Narechania, S. Singh, A. Scott, and A. Paasch for advice on methods and access to cluster computing resources.

## Notes

### Competing Interest Statement

The authors have declared no competing interest.

### Summary of Updates

Abstract, figures, and synteny results revised for greater clarity; Supplemental file 1 updated to include additional information.

## REFERENCES

1. Renvoisé E, Michon F. An Evo-Devo perspective on ever-growing teeth in mammals and dental stem cell maintenance. Front Physiol. 2014;5(324):1–12.

2. Tapaltsyan V, Eronen JT, Lawing AM, Sharir A, Janis C, Jernvall J, et al. Continuously growing rodent molars result from a predictable quantitative evolutionary change over 50 million years. Cell Rep. 2015;11(5):673–80.

3. LeBlanc ARH, Brink KS, Whitney MR, Abdala F, Reisz RR. Dental ontogeny in extinct synapsids reveals a complex evolutionary history of the mammalian tooth attachment system. Proc R Soc B Biol Sci. 2018 Nov 7;285(1890):20181792.

4. Saffar JL, Lasfargues JJ, Cherruau M. Alveolar bone and the alveolar process: the socket that is never stable. Periodontol 2000. 1997;13(1):76–90.

5. Davit-Béal T, Tucker AS, Sire JY. Loss of teeth and enamel in tetrapods: Fossil record, genetic data and morphological adaptations. J Anat. 2009;214(4):477–501.

6. Damuth J, Janis CM. On the relationship between hypsodonty and feeding ecology in ungulate mammals, and its utility in palaeoecology. Biol Rev. 2011;86(3):733–58.

7. Miletich I, Sharpe PT. Normal and abnormal dental development. Hum Mol Genet. 2003 Apr 2;12(suppl_1):R69–73.

8. Harada H, Kettunen P, Jung HS, Mustonen T, Wang YA, Thesleff I. Localization of putative stem cells in dental epithelium and their association with Notch and FGF signaling. J Cell Biol. 1999;147(1):105–20.

9. Tummers M, Thesleff I. Root or crown: a developmental choice orchestrated by the differential regulation of the epithelial stem cell niche in the tooth of two rodent species. Development. 2003;130(6):1049–57.

10. Thesleff I, Tummers M. Tooth organogenesis and regeneration. In: StemBook. Cambridge, MA: Harvard Stem Cell Institute; 2008.

11. Krivanek J, Buchtova M, Fried K, Adameyko I. Plasticity of dental cell types in development, regeneration, and evolution. J Dent Res. 2023 Jun 1;102(6):589–98.

12. Luan X, Ito Y, Diekwisch TGH. Evolution and development of Hertwig’s epithelial root sheath. Dev Dyn. 2006;235(5):1167–80.

13. Kumakami-Sakano M, Otsu K, Fujiwara N, Harada H. Regulatory mechanisms of Hertwig’s epithelial root sheath formation and anomaly correlated with root length. Exp Cell Res. 2014;325(2):78–82.

14. Wen Q, Jing J, Han X, Feng J, Yuan Y, Ma Y, et al. *Runx2* regulates mouse tooth root development via activation of WNT inhibitor *NOTUM*. J Bone Miner Res. 2020;35(11):2252–64.

15. Yang S, Choi H, Kim TH, Jeong JK, Liu Y, Harada H, et al. Cell dynamics in Hertwig’s epithelial root sheath are regulated by β-catenin activity during tooth root development. J Cell Physiol. 2021;236(7):5387–98.

16. Yamashiro T, Tummers M, Thesleff I. Expression of bone morphogenetic proteins and Msx genes during root formation. J Dent Res. 2003;82(3):172–6.

17. Yokohama-Tamaki T, Ohshima H, Fujiwara N, Takada Y, Ichimori Y, Wakisaka S, et al. Cessation of Fgf10 signaling, resulting in a defective dental epithelial stem cell compartment, leads to the transition from crown to root formation. Development. 2006;133(7):1359–66.

18. Ota MS, Vivatbutsin P, Nakahara T, Eto K. Tooth root development and the cell-based regenerative therapy. J Oral Tissue Eng. 2007;4(3):137–42.

19. Jernvall J, Thesleff I. Reiterative signaling and patterning during mammalian tooth morphogenesis. Mech Dev. 2000;92:19–29.

20. Harada H, Toyono T, Toyoshima K, Yamasaki M, Itoh N, Kato S, et al. FGF10 maintains stem cell compartment in developing mouse incisors. Dev Camb Engl. 2002;129(6):1533–41.

21. Tapaltsyan V, Charles C, Hu J, Mindell D, Ahituv N, Wilson GM, et al. Identification of novel *Fgf* enhancers and their role in dental evolution. Evol Dev. 2016;18(1):31–40.

22. Christensen MM, Hallikas O, Das Roy R, Väänänen V, Stenberg OE, Häkkinen TJ, et al. The developmental basis for scaling of mammalian tooth size. Proc Natl Acad Sci. 2023 Jun 20;120(25):e2300374120.

23. Chen ZJ. Genetic and epigenetic mechanisms for gene expression and phenotypic cariation in plant polyploids. Annu Rev Plant Biol. 2007;58(1):377–406.

24. Stranger BE, Forrest MS, Dunning M, Ingle CE, Beazley C, Thorne N, et al. Relative impact of nucleotide and copy number variation on gene expression phenotypes. Science. 2007 Feb 9;315(5813):848–53.

25. Romero IG, Ruvinsky I, Gilad Y. Comparative studies of gene expression and the evolution of gene regulation. Nat Rev Genet. 2012 Jul;13(7):505–16.

26. de Montaigu A, Giakountis A, Rubin M, Tóth R, Cremer F, Sokolova V, et al. Natural diversity in daily rhythms of gene expression contributes to phenotypic variation. Proc Natl Acad Sci. 2015 Jan 20;112(3):905–10.

27. Erwin DH, Davidson EH. The last common bilaterian ancestor. Development. 2002 Jul 1;129(13):3021–32.

28. Irie N, Kuratani S. Comparative transcriptome analysis reveals vertebrate phylotypic period during organogenesis. Nat Commun. 2011;2:248.

29. Koonin EV. Evolution of genome architecture. Int J Biochem Cell Biol. 2009 Feb 1;41(2):298–306.

30. Wray GA. The evolutionary significance of cis-regulatory mutations. Nat Rev Genet. 2007 Mar;8(3):206–16.

31. Acemel RD, Maeso I, Gómez-Skarmeta JL. Topologically associated domains: a successful scaffold for the evolution of gene regulation in animals. WIREs Dev Biol. 2017;6(3):e265.

32. Coghlan A, Eichler EE, Oliver SG, Paterson AH, Stein L. Chromosome evolution in eukaryotes: a multi-kingdom perspective. Trends Genet. 2005 Dec 1;21(12):673–82.

33. Swenson KM, Blanchette M. Large-scale mammalian genome rearrangements coincide with chromatin interactions. Bioinformatics. 2019 Jul 15;35(14):i117–26.

34. Long HS, Greenaway S, Powell G, Mallon AM, Lindgren CM, Simon MM. Making sense of the linear genome, gene function and TADs. Epigenetics Chromatin. 2022 Jan 29;15(1):4.

35. Harmston N, Ing-Simmons E, Tan G, Perry M, Merkenschlager M, Lenhard B. Topologically associating domains are ancient features that coincide with Metazoan clusters of extreme noncoding conservation. Nat Commun. 2017 Sep 5;8(1):441.

36. Szabo Q, Bantignies F, Cavalli G. Principles of genome folding into topologically associating domains. Sci Adv. 2019 Apr 10;5(4):eaaw1668.

37. Das Roy R, Hallikas O, Christensen MM, Renvoisé E, Jernvall J. Chromosomal neighbourhoods allow identification of organ specific changes in gene expression. PLOS Comput Biol. 2021 Sep 10;17(9):e1008947.

38. Torelli F, Zander S, Ellerbrok H, Kochs G, Ulrich RG, Klotz C, et al. Recombinant IFN-γ from the bank vole *Myodes glareolus*: a novel tool for research on rodent reservoirs of zoonotic pathogens. Sci Rep. 2018;8(1):1–11.

39. Kloch A, Babik W, Bajer A, Siński E, Radwan J. Effects of an MHC-DRB genotype and allele number on the load of gut parasites in the bank vole *Myodes glareolus*. Mol Ecol. 2010;19(SUPPL. 1):255–65.

40. Migalska M, Sebastian A, Konczal M, Kotlík P, Radwan J. *De novo* transcriptome assembly facilitates characterisation of fast-evolving gene families, MHC class I in the bank vole (*Myodes glareolus)*. Heredity. 2017;118(4):348–57.

41. Appleton J, Lee KM, Sawicka Kapusta K, Damek M, Cooke M. The heavy metal content of the teeth of the bank vole (*Clethrionomys glareolus*) as an exposure marker of environmental pollution in Poland. Environ Pollut. 2000;110:441–9.

42. Gdula-Argasińska J, Appleton J, Sawicka-Kapusta K, Spence B. Further investigation of the heavy metal content of the teeth of the bank vole as an exposure indicator of environmental pollution in Poland. Environ Pollut. 2004;131(1):71–9.

43. Hallikas O, Das Roy R, Christensen MM, Renvoisé E, Sulic AM, Jernvall J. System-level analyses of keystone genes required for mammalian tooth development. J Exp Zoolog B Mol Dev Evol. 2021;336(1):7–17.

44. Yang Z. PAML 4: Phylogenetic analysis by maximum likelihood. Mol Biol Evol. 2007 Aug 1;24(8):1586–91.

45. Zhang J, Nielsen R, Yang Z. Evaluation of an improved branch-site likelihood method for detecting positive selection at the molecular level. Mol Biol Evol. 2005 Dec;22(12):2472–9.

46. Yang Z, Wong WSW, Nielsen R. Bayes Empirical Bayes inference of amino acid sites under positive selection. Mol Biol Evol. 2005 Apr 1;22(4):1107–18.

47. Drummond DA, Bloom JD, Adami C, Wilke CO, Arnold FH. Why highly expressed proteins evolve slowly. Proc Natl Acad Sci. 2005 Oct 4;102(40):14338–43.

48. Kosiol C, Vinař T, Fonseca RR da, Hubisz MJ, Bustamante CD, Nielsen R, et al. Patterns of positive selection in six mammalian genomes. PLOS Genet. 2008 Aug 1;4(8):e1000144.

49. Martincorena I, Luscombe NM. Non-random mutation: The evolution of targeted hypermutation and hypomutation. BioEssays. 2013;35(2):123–30.

50. Martincorena I, Roshan A, Gerstung M, Ellis P, Van Loo P, McLaren S, et al. High burden and pervasive positive selection of somatic mutations in normal human skin. Science. 2015 May 22;348(6237):880–6.

51. Keränen SVE, Åberg T, Kettunen P, Thesleff I, Jernvall J. Association of developmental regulatory genes with the development of different molar tooth shapes in two species of rodents. Dev Genes Evol. 1998;208(9):477–86.

52. Jernvall J, Keränen SVE, Thesleff I. Evolutionary modification of development in mammalian teeth: Quantifying gene expression patterns and topography. Proc Natl Acad Sci. 2000;97(26):14444–8.

53. Hughes AL. The evolution of functionally novel proteins after gene duplication. Proc R Soc Lond B Biol Sci. 1997 Jan;256(1346):119–24.

54. Wagner A. Selection and gene duplication: a view from the genome. Genome Biol. 2002 Apr 15;3(5):reviews1012.1.

55. David KT, Oaks JR, Halanych KM. Patterns of gene evolution following duplications and speciations in vertebrates. PeerJ. 2020 Mar 31;8:e8813.

56. Copley SD. Evolution of new enzymes by gene duplication and divergence. FEBS J. 2020;287(7):1262–83.

57. Fisher LW. DMP1 and DSPP: Evidence for duplication and convergent evolution of two SIBLING proteins. Cells Tissues Organs. 2011 Aug;194(2–4):113–8.

58. Bouleftour W, Juignet L, Bouet G, Granito RN, Vanden-Bossche A, Laroche N, et al. The role of the SIBLING, Bone Sialoprotein in skeletal biology — Contribution of mouse experimental genetics. Matrix Biol. 2016 May 1;52–54:60–77.

59. Felszeghy S, Módis L, Németh P, Nagy G, Zelles T, Agre P, et al. Expression of aquaporin isoforms during human and mouse tooth development. Arch Oral Biol. 2004 Apr 1;49(4):247–57.

60. Yoshii T, Harada F, Saito I, Nozawa-Inoue K, Kawano Y, Maeda T. Immunoexpression of aquaporin-1 in the rat periodontal ligament during experimental tooth movement. Biomed Res. 2012;33(4):225–33.

61. Zhang X, Zhao J, Li C, Gao S, Qiu C, Liu P, et al. *DSPP* mutation in dentinogenesis imperfecta Shields type II. Nat Genet. 2001 Feb;27(2):151–2.

62. de La Dure-Molla M, Philippe Fournier B, Berdal A. Isolated dentinogenesis imperfecta and dentin dysplasia: revision of the classification. Eur J Hum Genet. 2015 Apr;23(4):445–51.

63. Shields ED, Bixler D, El-Kafrawy AM. A proposed classification for heritable human dentine defects with a description of a new entity. Arch Oral Biol. 1973 Apr 1;18(4):543–IN7.

64. Sreenath T, Thyagarajan T, Hall B, Longenecker G, D’Souza R, Hong S, et al. Dentin Sialophosphoprotein knockout mouse teeth display widened predentin zone and develop defective dentin mineralization similar to human dentinogenesis imperfecta type III. J Biol Chem. 2003 Jul 4;278(27):24874–80.

65. Verdelis K, Ling Y, Sreenath T, Haruyama N, MacDougall M, van der Meulen MCH, et al. DSPP effects on *in vivo* bone mineralization. Bone. 2008 Dec 1;43(6):983–90.

66. Chen Y, Zhang Y, Ramachandran A, George A. DSPP is essential for normal development of the dental-craniofacial complex. J Dent Res. 2016 Mar 1;95(3):302–10.

67. von Marschall Z, Mok S, Phillips MD, McKnight DA, Fisher LW. Rough endoplasmic reticulum trafficking errors by different classes of mutant dentin sialophosphoprotein (DSPP) cause dominant negative effects in both dentinogenesis imperfecta and dentin dysplasia by entrapping normal DSPP. J Bone Miner Res. 2012;27(6):1309–21.

68. Smith BL, Preston GM, Spring FA, Anstee DJ, Agre P. Human red cell aquaporin CHIP. I. Molecular characterization of ABH and Colton blood group antigens. J Clin Invest. 1994 Sep 1;94(3):1043–9.

69. Jordan IK, Mariño-Ramírez L, Koonin EV. Evolutionary significance of gene expression divergence. Gene. 2005 Jan 17;345(1):119–26.

70. Warnefors M, Kaessmann H. Evolution of the correlation between expression divergence and protein divergence in mammals. Genome Biol Evol. 2013;5(7):1324–35.

71. Jernvall J, Thesleff I. Tooth shape formation and tooth renewal: evolving with the same signals. Development. 2012;139(19):3487–97.

72. Mitsiadis TA. Role of Islet1 in the patterning of murine dentition. Development. 2003;130(18):4451–60.

73. Charles C, Pantalacci S, Peterkova R, Tafforeau P, Laudet V, Viriot L. Effect of *eda* loss of function on upper jugal tooth morphology. Anat Rec. 2009;292(2):299–308.

74. Zurowski C, Jamniczky H, Graf D, Theodor J. Deletion/loss of bone morphogenetic protein 7 changes tooth morphology and function in *Mus musculus*: implications for dental evolution in mammals. R Soc Open Sci. 2018 Jan 3;5(1):170761.

75. Cardoso-Moreira M, Halbert J, Valloton D, Velten B, Chen C, Shao Y, et al. Gene expression across mammalian organ development. Nature. 2019 Jul;571(7766):505–9.

76. Finarelli JA, Flynn JJ. Ancestral state reconstruction of body size in the Caniformia (Carnivora, Mammalia): the effects of incorporating data from the fossil record. Syst Biol. 2006;55(2):301–13.

77. Welker F, Collins MJ, Thomas JA, Wadsley M, Brace S, Cappellini E, et al. Ancient proteins resolve the evolutionary history of Darwin’s South American ungulates. Nature. 2015 Jun;522(7554):81–4.

78. Warinner C, Korzow Richter K, Collins MJ. Paleoproteomics. Chem Rev. 2022 Aug 24;122(16):13401–46.

79. Zheng GXY, Lau BT, Schnall-Levin M, Jarosz M, Bell JM, Hindson CM, et al. Haplotyping germline and cancer genomes with high-throughput linked-read sequencing. Nat Biotechnol. 2016 Feb;34:303.

80. Marks P, Garcia S, Martinez A, Belhocine K. Resolving the full spectrum of human genome variation using linked-reads. 2017;

81. Weisenfeld NI, Kumar V, Shah P, Church DM, Jaffe DB. Direct determination of diploid genome sequences. Genome Res. 2017;27(5):757–67.

82. Vurture GW, Sedlazeck FJ, Nattestad M, Underwood CJ, Fang H, Gurtowski J, et al. GenomeScope: fast reference-free genome profiling from short reads. Bioinformatics. 2017 Jul 15;33(14):2202–4.

83. Koren S, Walenz BP, Berlin K, Miller JR, Bergman NH, Phillippy AM. Canu: scalable and accurate long-read assembly via adaptive k-mer weighting and repeat separation. Genome Res. 2017 May 1;27(5):722–36.

84. Warren RL. RAILS and Cobbler: Scaffolding and automated finishing of draft genomes using long DNA sequences. J Open Source Softw. 2016 Nov 17;1(7):116.

85. Gurevich A, Saveliev V, Vyahhi N, Tesler G. QUAST: quality assessment tool for genome assemblies. Bioinformatics. 2013 Apr 15;29(8):1072–5.

86. Simão FA, Waterhouse RM, Ioannidis P, Kriventseva EV, Zdobnov EM. BUSCO: assessing genome assembly and annotation completeness with single-copy orthologs. Bioinformatics. 2015 Oct 1;31(19):3210–2.

87. Cantarel BL, Korf I, Robb SMC, Parra G, Ross E, Moore B, et al. MAKER: an easy-to-use annotation pipeline designed for emerging model organism genomes. Genome Res. 2008;18:188–96.

88. Campbell MS, Law M, Holt C, Stein JC, Moghe GD, Hufnagel DE, et al. MAKER-P: A tool kit for the rapid creation, management, and quality control of plant genome annotations. Plant Physiol. 2014 Feb 1;164(2):513–24.

89. Campbell MS, Holt C, Moore B, Yandell M. Genome annotation and curation using MAKER and MAKER-P. Curr Protoc Bioinforma. 2014 Dec 12;48:4.11.1–4.11.39.

90. Grabherr MG, Haas BJ, Yassour M, Levin JZ, Thompson DA, Amit I, et al. Trinity: reconstructing a full-length transcriptome without a genome from RNA-Seq data. Nat Biotechnol. 2011 May 15;29(7):644–52.

91. Wheeler TJ, Clements J, Eddy SR, Hubley R, Jones TA, Jurka J, et al. Dfam: a database of repetitive DNA based on profile hidden Markov models. Nucleic Acids Res. 2013 Jan;41(Database issue):D70–82.

92. Caballero J, Smit AFA, Hood L, Glusman G. Realistic artificial DNA sequences as negative controls for computational genomics. Nucleic Acids Res. 2014 Jul;42(12):e99.

93. Hubley R, Finn RD, Clements J, Eddy SR, Jones TA, Bao W, et al. The Dfam database of repetitive DNA families. Nucleic Acids Res. 2016 Jan 4;44(D1):D81–9.

94. Hu J, Zheng Y, Shang X. MiteFinder: A fast approach to identify miniature inverted-repeat transposable elements on a genome-wide scale. In: 2017 IEEE International Conference on Bioinformatics and Biomedicine (BIBM). 2017. p. 164–8.

95. Gremme G, Steinbiss S, Kurtz S. GenomeTools: A comprehensive software library for efficient processing of structured genome annotations. IEEE/ACM Trans Comput Biol Bioinform. 2013 May 1;10(03):645–56.

96. Smit A, Hubley R. RepeatModeler Open-1.0. 2008.

97. Keller O, Kollmar M, Stanke M, Waack S. A novel hybrid gene prediction method employing protein multiple sequence alignments. Bioinformatics. 2011 Mar 15;27(6):757–63.

98. Korf I. Gene finding in novel genomes. BMC Bioinformatics. 2004 May 14;5(1):59.

99. Campbell MS. compare_annotations_3.2.pl [Internet]. 2015. Available from: https://github.com/mscampbell/Genome_annotation/blob/master/compare_annotations_3.2.pl

100. Eilbeck K, Moore B, Holt C, Yandell M. Quantitative measures for the management and comparison of annotated genomes. BMC Bioinformatics. 2009 Feb 23;10(1):67.

101. Liu D, Hunt M, Tsai IJ. Inferring synteny between genome assemblies: a systematic evaluation. BMC Bioinformatics. 2018 Jan;19(1):26.

102. Emms DM, Kelly S. OrthoFinder: solving fundamental biases in whole genome comparisons dramatically improves orthogroup inference accuracy. Genome Biol. 2015 Aug 6;16(1):157.

103. Katoh K, Misawa K, Kuma K, Miyata T. MAFFT: a novel method for rapid multiple sequence alignment based on fast Fourier transform. Nucleic Acids Res. 2002 Jul;30(14):3059–66.

104. Lefort V, Desper R, Gascuel O. FastME 2.0: A comprehensive, accurate, and fast distance-based phylogeny inference program. Mol Biol Evol. 2015 Oct 1;32(10):2798–800.

105. Farrer RA. Synima: a Synteny imaging tool for annotated genome assemblies. BMC Bioinformatics. 2017 Nov 21;18(1):507.

106. Wang Y, Tang H, DeBarry JD, Tan X, Li J, Wang X, et al. MCScanX: a toolkit for detection and evolutionary analysis of gene synteny and collinearity. Nucleic Acids Res. 2012 Apr;40(7):e49.

107. Zhao T, Schranz ME. Network approaches for plant phylogenomic synteny analysis. Curr Opin Plant Biol. 2017 Apr 1;36:129–34.

108. Zhao T, Holmer R, de Bruijn S, Angenent GC, van den Burg HA, Schranz ME. Phylogenomic synteny network analysis of MADS-Box transcription factor genes reveals lineage-specific transpositions, ancient tandem duplications, and deep positional conservation. Plant Cell. 2017 Jun 1;29(6):1278–92.

109. Zhao T, Schranz ME. Network-based microsynteny analysis identifies major differences and genomic outliers in mammalian and angiosperm genomes. Proc Natl Acad Sci. 2019 Feb 5;116(6):2165–74.

110. Sievers F, Higgins DG. Clustal Omega. Curr Protoc Bioinforma. 2014;48(1):3.13.1–3.13.16.

111. Suyama M, Torrents D, Bork P. PAL2NAL: robust conversion of protein sequence alignments into the corresponding codon alignments. Nucleic Acids Res. 2006 Jul 1;34(suppl_2):W609–12.

112. Wong WSW, Yang Z, Goldman N, Nielsen R. Accuracy and power of statistical methods for detecting adaptive evolution in protein coding sequences and for identifying positively selected sites. Genetics. 2004 Oct 1;168(2):1041–51.

113. Löytynoja A, Vilella AJ, Goldman N. Accurate extension of multiple sequence alignments using a phylogeny-aware graph algorithm. Bioinformatics. 2012 Jul 1;28(13):1684–91.

114. Jones DT. Protein secondary structure prediction based on position-specific scoring matrices. J Mol Biol. 1999 Sep 17;292(2):195–202.

115. Buchan DWA, Jones DT. The PSIPRED protein analysis workbench: 20 years on. Nucleic Acids Res. 2019 Jul 2;47(W1):W402–7.

116. Ewels PA, Peltzer A, Fillinger S, Patel H, Alneberg J, Wilm A, et al. The nf-core framework for community-curated bioinformatics pipelines. Nat Biotechnol. 2020 Mar;38(3):276–8.

117. Andrews S. FastQC: a quality control tool for high throughput sequence data. [Internet]. 2010. Available from: http://www.bioinformatics.babraham.ac.uk/projects/fastqc

118. Martin M. Cutadapt removes adapter sequences from high-throughput sequencing reads. EMBnet.journal. 2011 May 2;17(1):10–2.

119. Kopylova E, Noé L, Touzet H. SortMeRNA: fast and accurate filtering of ribosomal RNAs in metatranscriptomic data. Bioinformatics. 2012 Dec 1;28(24):3211–7.

120. Patro R, Duggal G, Love MI, Irizarry RA, Kingsford C. Salmon provides fast and bias-aware quantification of transcript expression. Nat Methods. 2017 Apr;14(4):417–9.

121. Love MI, Huber W, Anders S. Moderated estimation of fold change and dispersion for RNA-seq data with DESeq2. Genome Biol. 2014 Dec 5;15(12):550.

122. Lindblad-Toh K, Garber M, Zuk O, Lin MF, Parker BJ, Washietl S, et al. A high-resolution map of human evolutionary constraint using 29 mammals. Nature. 2011 Oct;478(7370):476–82.

123. Weyrich A, Schüllermann T, Heeger F, Jeschek M, Mazzoni CJ, Chen W, et al. Whole genome sequencing and methylome analysis of the wild guinea pig. BMC Genomics. 2014 Nov 28;15(1):1036.

124. Gossmann TI, Ralser M. Marmota marmota. Trends Genet. 2020 May;36(5):383–4.

125. Di Palma F, Alföldi J, Johnson J, Berlin A, Gnerre S, Jaffe D, et al. The draft genome of *Microtus ochrogaster*. Broad Inst [Internet]. 2012; Available from: https://www.ncbi.nlm.nih.gov/bioproject/72443

126. Mouse Genome Sequencing Consortium, Waterston RH, Lindblad-Toh K, Birney E, Rogers J, Abril JF, et al. Initial sequencing and comparative analysis of the mouse genome. Nature. 2002 Dec 5;420(6915):520–62.

127. Di Palma F, Alföldi J, Johnson J, Berlin A, Gnerre S, Jaffe D, et al. The draft genome of *Jaculus jaculus*. Broad Inst [Internet]. 2012; Available from: https://www.ncbi.nlm.nih.gov/bioproject/72445

128. Gibbs RA, Weinstock GM, Metzker ML, Muzny DM, Sodergren EJ, Scherer S, et al. Genome sequence of the Brown Norway rat yields insights into mammalian evolution. Nature. 2004 Apr;428(6982):493–521.

129. Kolmogorov M, Armstrong J, Raney BJ, Streeter I, Dunn M, Yang F, et al. Chromosome assembly of large and complex genomes using multiple references. Genome Res. 2018 Nov 1;28(11):1720–32.

130. Lilue J, Doran AG, Fiddes IT, Abrudan M, Armstrong J, Bennett R, et al. Sixteen diverse laboratory mouse reference genomes define strain-specific haplotypes and novel functional loci. Nat Genet. 2018 Nov;50(11):1574–83.

131. Couger MB, Arévalo L, Campbell P. A high quality genome for *Mus spicilegus*, a close relative of house mice with unique social and ecological adaptations. G3 GenesGenomesGenetics. 2018 May 24;8(7):2145–52.

132. Chinese hamster CHOK1GS assembly and gene annotation. Horiz Eagle [Internet]. 2017; Available from: https://www.ensembl.org/Cricetulus_griseus_chok1gshd/Info/Annotation

133. Di Palma F, Alföldi J, Johnson J, Berlin A, Gnerre S, Jaffe D, et al. The draft genome of *Mesocricetus auratus*. Broad Inst [Internet]. 2012; Available from: https://www.ncbi.nlm.nih.gov/bioproject/77669

134. Lassance JM, Hopi Hoekstra. Improved assembly of the deer mouse *Peromyscus maniculatus* genome. Harv Univ Hughes Med Inst [Internet]. 2018; Available from: https://www.ncbi.nlm.nih.gov/bioproject/494228

135. Fang X, Nevo E, Han L, Levanon EY, Zhao J, Avivi A, et al. Genome-wide adaptive complexes to underground stresses in blind mole rats *Spalax*. Nat Commun. 2014 Jun 3;5(1):3966.

136. Di Palma F, Alföldi J, Johnson J, Berlin A, Gnerre S, Jaffe D, et al. The draft genome of *Octodon degu*. Broad Inst [Internet]. 2012; Available from: https://www.ncbi.nlm.nih.gov/bioproject/74595

137. Keane M, Craig T, Alföldi J, Berlin AM, Johnson J, Seluanov A, et al. The Naked Mole Rat Genome Resource: facilitating analyses of cancer and longevity-related adaptations. Bioinforma Oxf Engl. 2014 Dec 15;30(24):3558–60.

138. Di Palma F, Alföldi J, Johnson J, Berlin A, Gnerre S, Jaffe D, et al. The draft genome of *Chinchilla lanigera*. Broad Inst [Internet]. 2012; Available from: https://www.ncbi.nlm.nih.gov/bioproject/68239

139. V. Federov, Dalen L, Olsen RA, Goropashnaya AV, Barnes BM. The genome of the Arctic ground squirrel *Urocitellus parryii*. Inst Arct Biol [Internet]. 2018; Available from: https://www.ncbi.nlm.nih.gov/bioproject/477386

140. Di Palma F, Alföldi J, Johnson J, Berlin A, Gnerre S, Jaffe D, et al. The draft genome of *Ictidomys tridecemlineatus*. Broad Inst [Internet]. 2012; Available from: https://www.ncbi.nlm.nih.gov/bioproject/61725

141. Schneider VA, Graves-Lindsay T, Howe K, Bouk N, Chen HC, Kitts PA, et al. Evaluation of GRCh38 and de novo haploid genome assemblies demonstrates the enduring quality of the reference assembly. Genome Res. 2017 May 1;27(5):849–64.

142. Csárdi G, Nepusz T, Traag V, Horvát S, Zanini F, Noom D, et al. igraph: Network analysis and visualization in R [Internet]. 2024. Available from: https://CRAN.R-project.org/package=igraph

