## Additional file 4: Structure predictions for "Vole genomics links determinate and indeterminate growth of teeth"

**Additional file 4 – Structure predictions**
PSIPRED secondary structure predictions for *Dspp* and *Aqp1*, including both extant species with unrooted molars and the hypothetical ancestral nodes where ancestral sequence reconstruction showed the amino acid at the site under positive selection had changed.

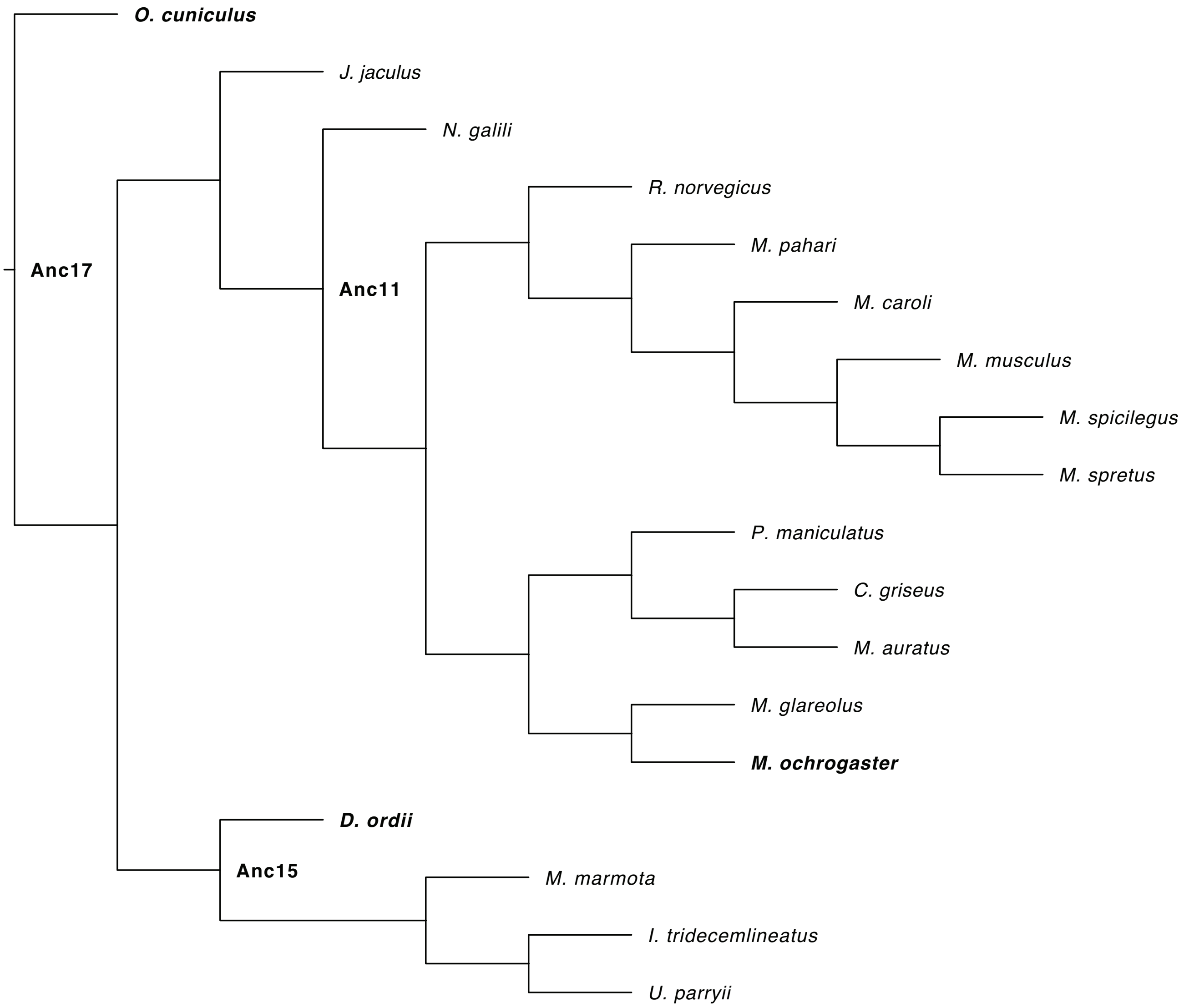


**Figure A4.1** – Gene tree for *Dspp* based on OrthoFinder analysis. Labeled internal nodes represent hypothetical ancestors where the amino acid at the site under branch-specific positive selection changed in the ancestral state reconstruction. Bold text indicates the internal nodes and tips for which we made secondary structure predictions.

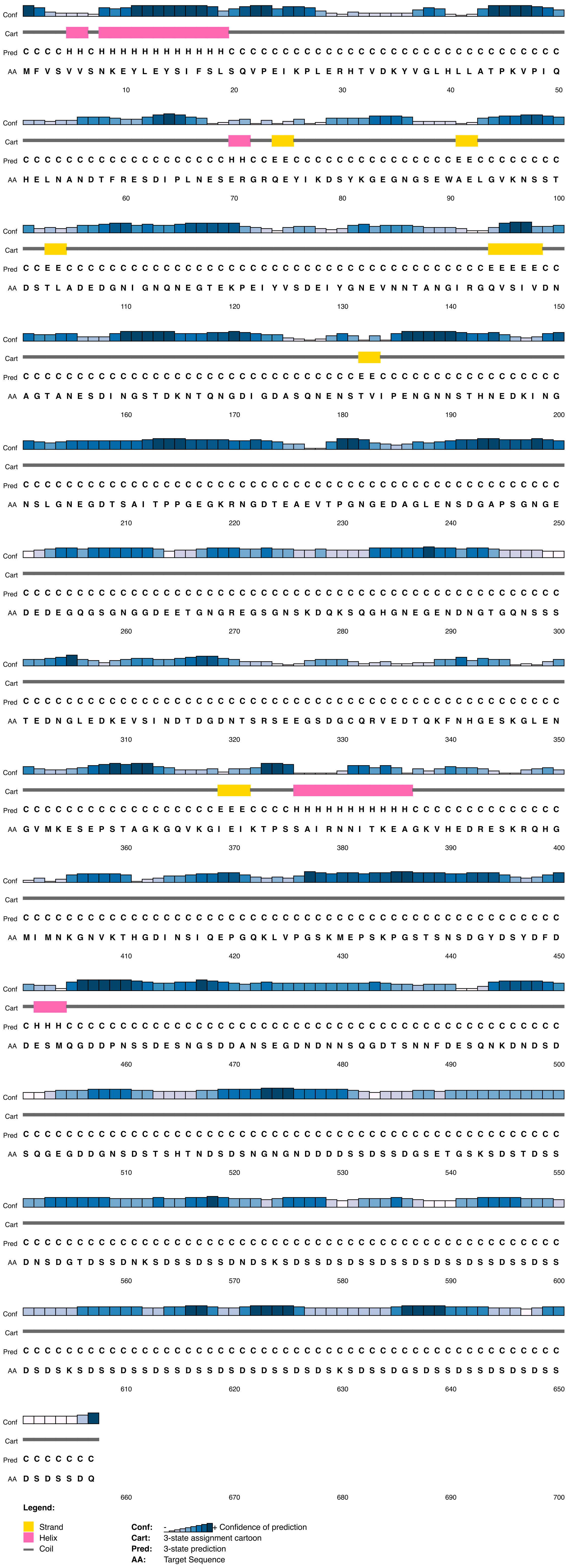


**Figure A4.2** – *Dspp* secondary structure prediction for *Oryctolagus cuniculus*. At the site under positive selection, *O. cuniculus* had a leucine and its ancestor, Anc17, had an asparagine (see Figure 6 in the main text).

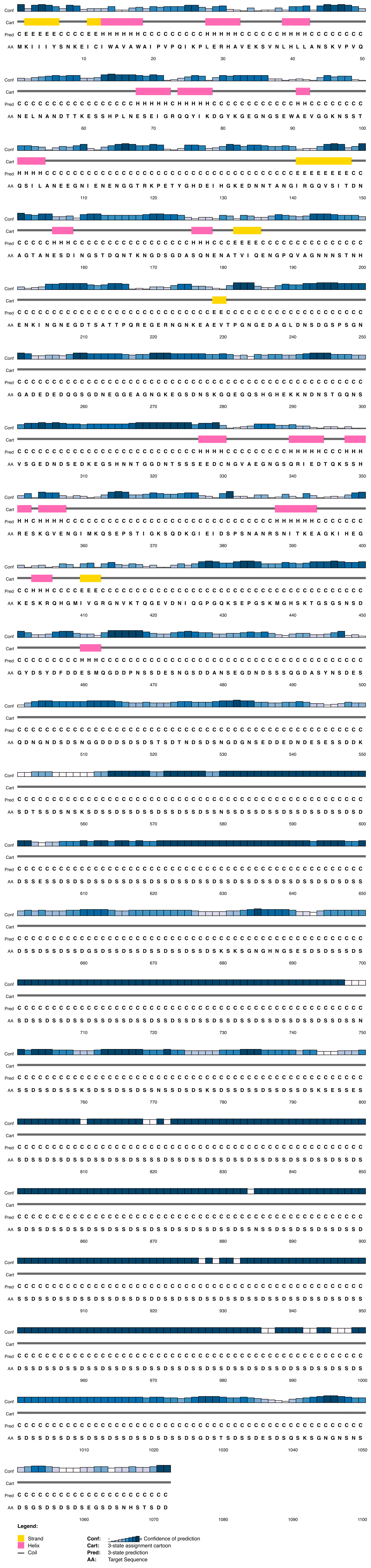


**Figure A4.3** – *Dspp* secondary structure prediction for Anc17, the ancestor of *Oryctolagus cuniculus*. Anc17 had an asparagine at the site under positive selection which was subsituted for a leucine in *O. cuniculus* (see Figure 6 in the main text).

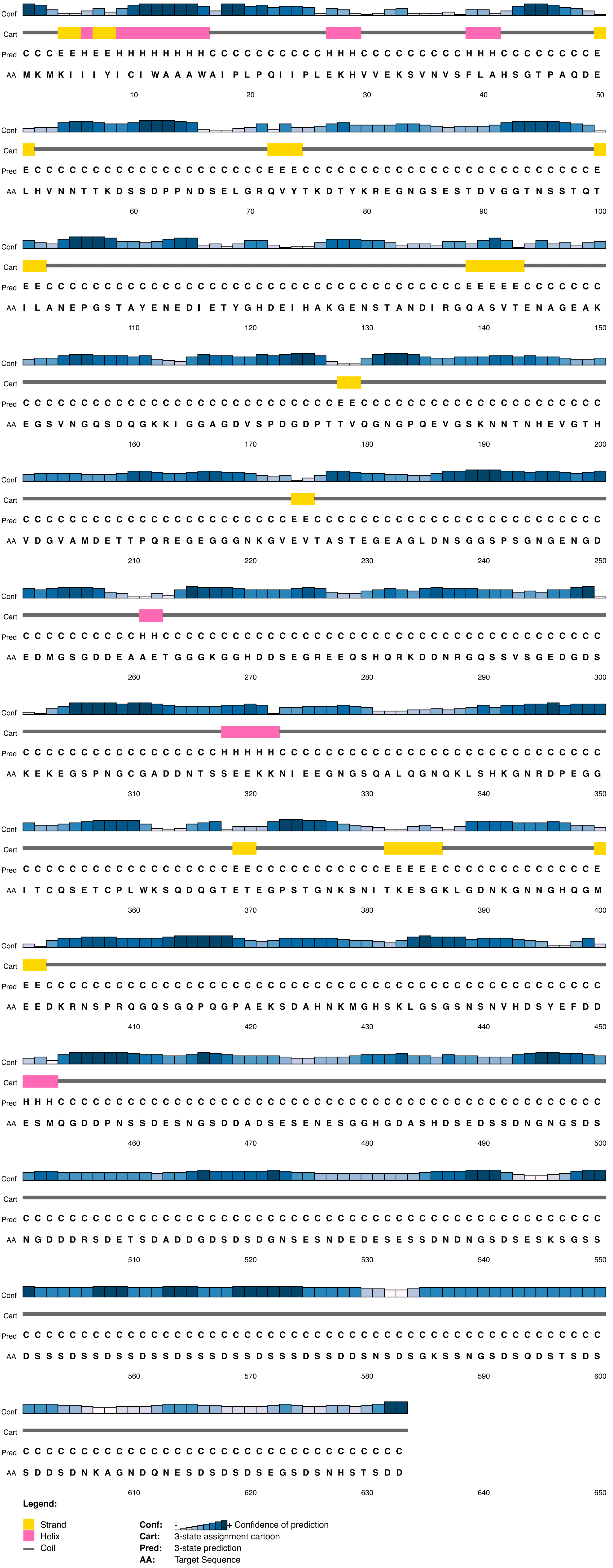


**Figure A4.4** – *Dspp* secondary structure prediction for *Microtus ochrogaster*. At the site under positive selection, *M. ochrogaster* had a histidine and its ancestor, Anc11, had an asparagine (see Figure 6 in the main text).

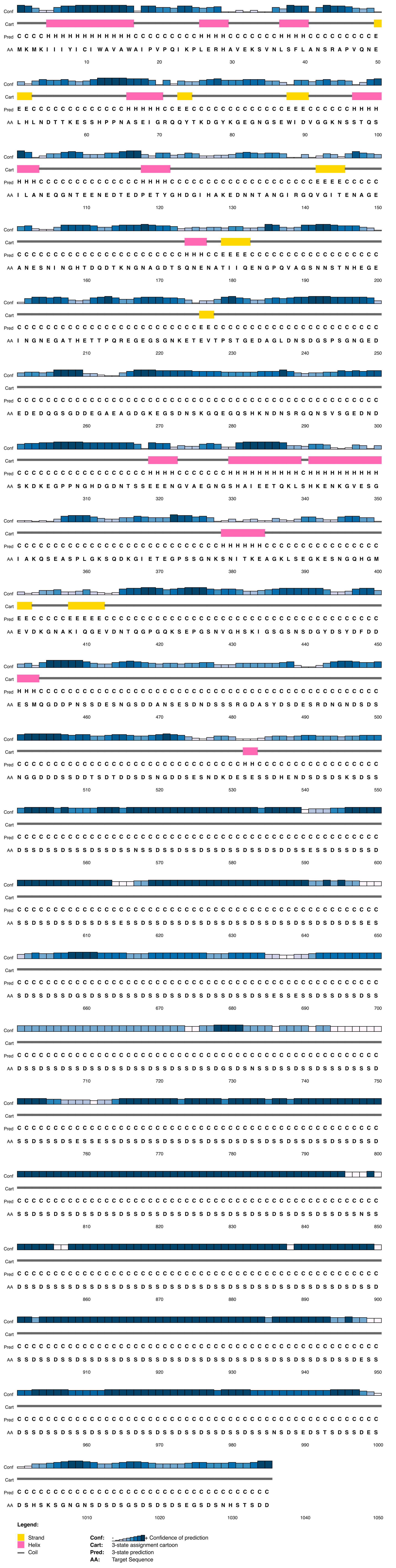


**Figure A4.5** – *Dspp* secondary structure prediction for Anc11, the ancestor of *Microtus ochrogaster*. Anc11 had an asparagine at the site under positive selection which was subsituted for a histidine in *M. ochrogaster* (see Figure 6 in the main text).

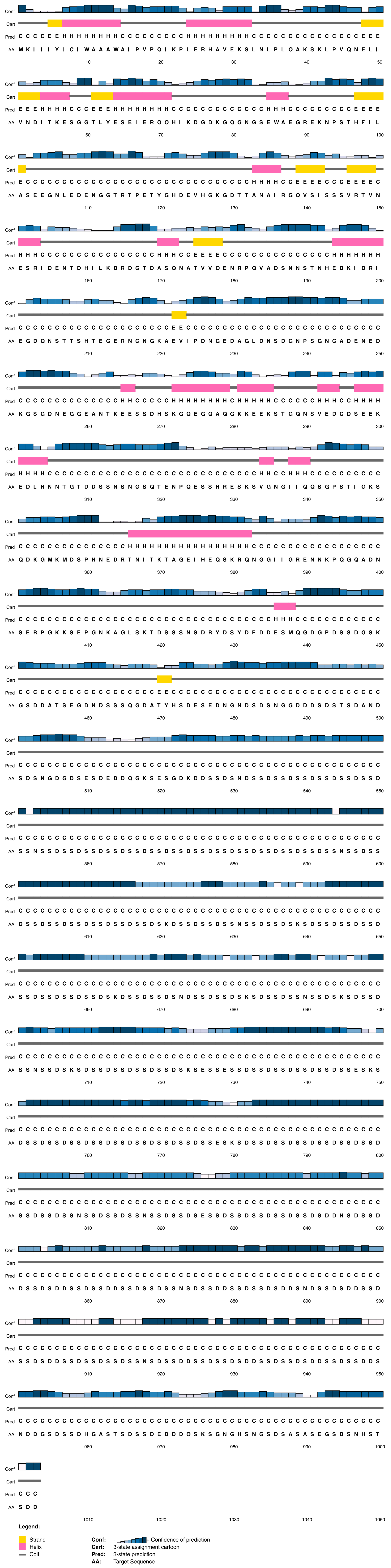


**Figure A4.6** – *Dspp* secondary structure prediction for *Dipodomys ordii*. At the site under positive selection, *D. ordii* had an aspartic acid and its ancestor, Anc15, had an asparagine (see Figure 6 in the main text).

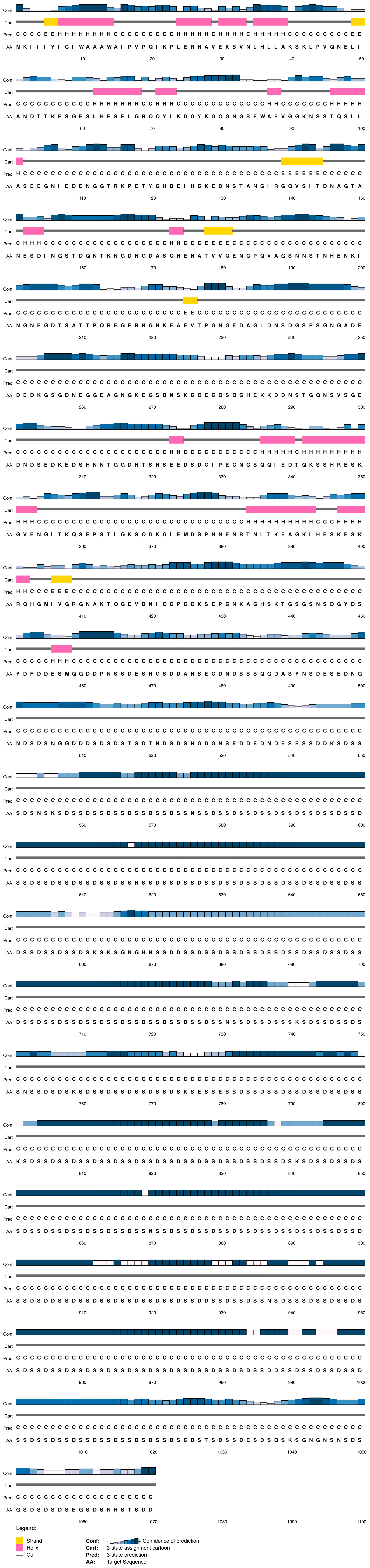


**Figure A4.7** – *Dspp* secondary structure prediction for Anc15, the ancestor of *Dipodomys ordii*. Anc15 had an asparagine at the site under positive selection which was subsituted for an aspartic acid in *D. ordii* (see Figure 6 in the main text).

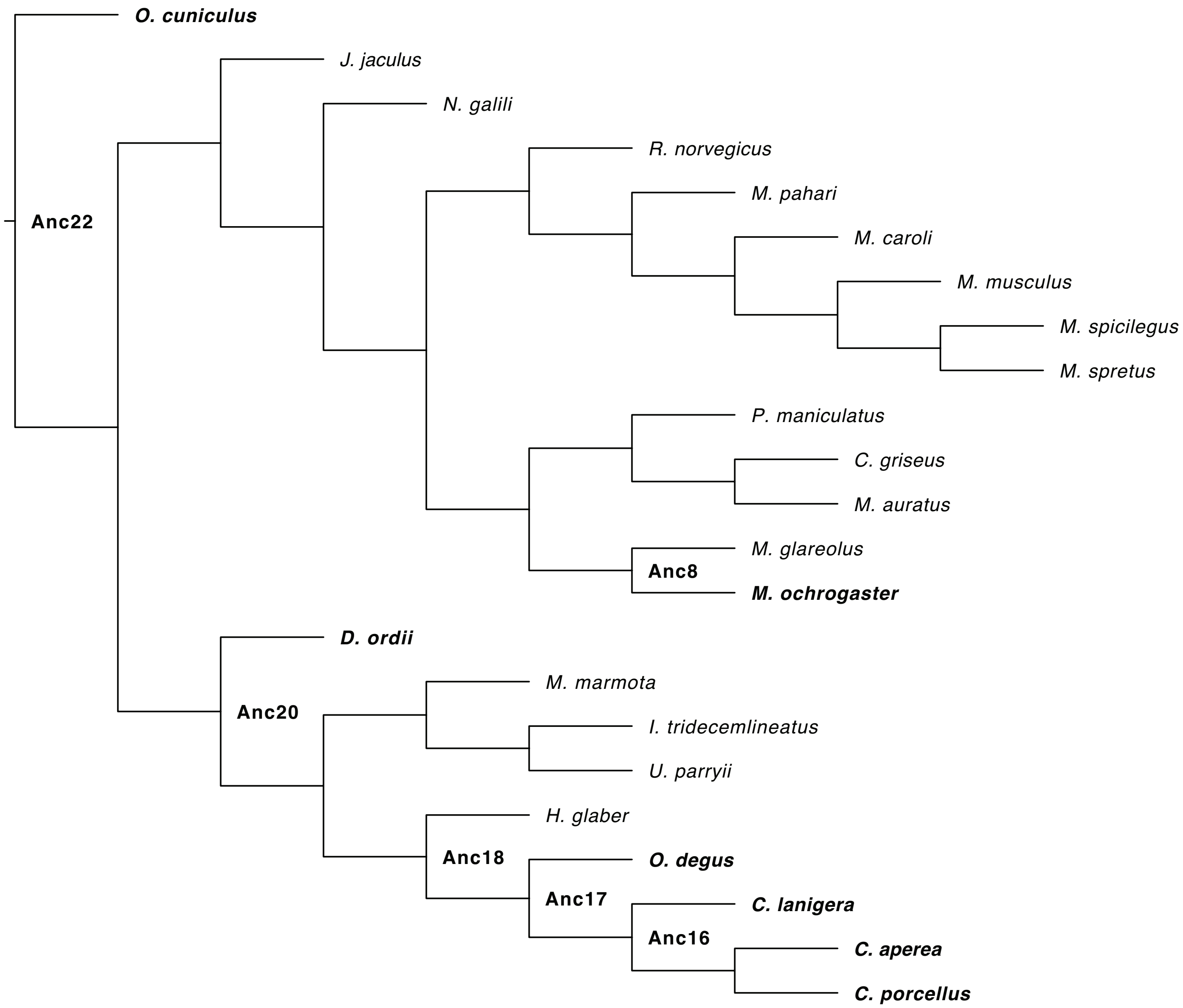


**Figure A4.8** – Gene tree for *Aqp1* based on OrthoFinder analysis. Labeled internal nodes represent hypothetical ancestors where the amino acid at the site under branch-specific positive selection changed in the ancestral state reconstruction. Bold text indicates the internal nodes and tips for which we made secondary structure predictions.

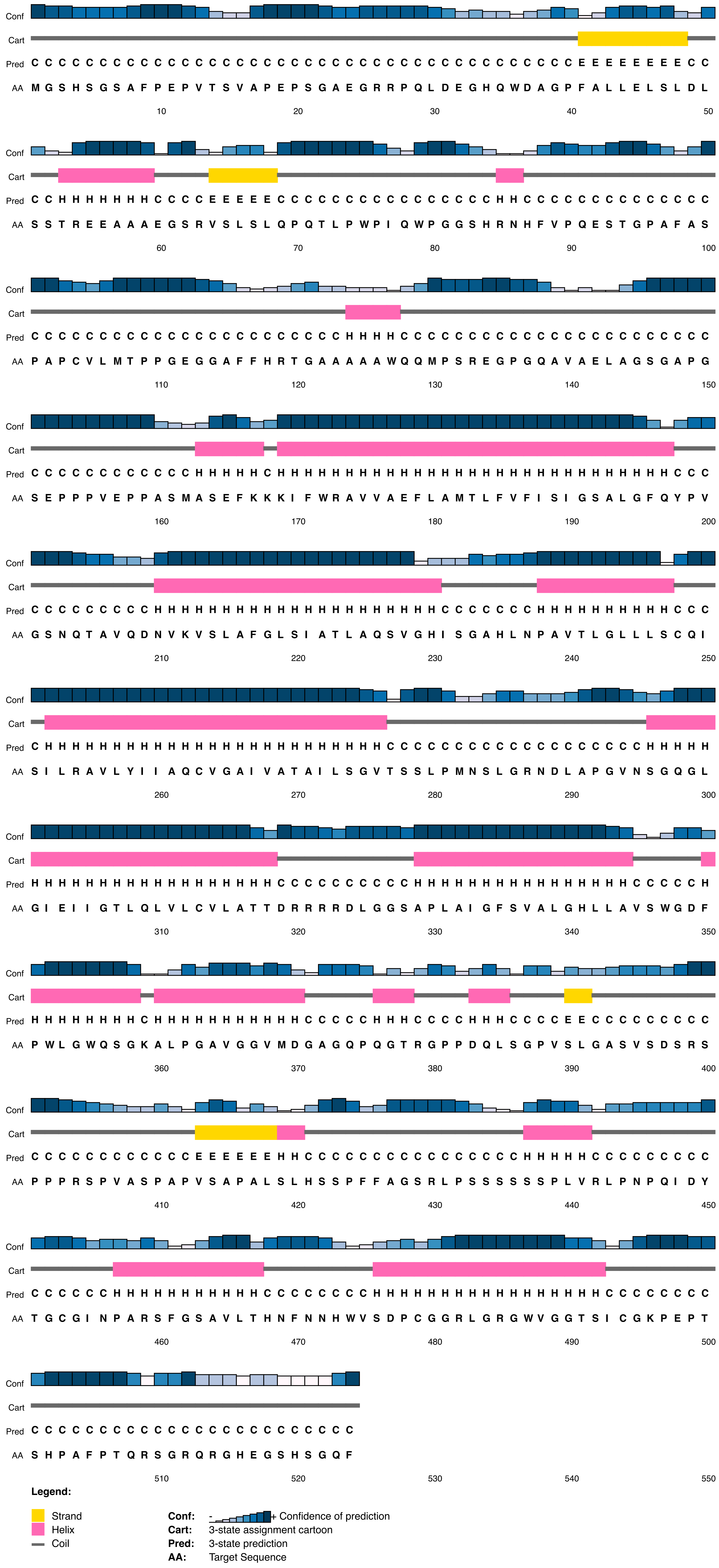


**Figure A4.9** – *Aqp1* secondary structure prediction for *Oryctolagus cuniculus*. At the site under positive selection, *O. cuniculus* and its ancestor, Anc22, both had a proline (see Figure 6 in the main text).

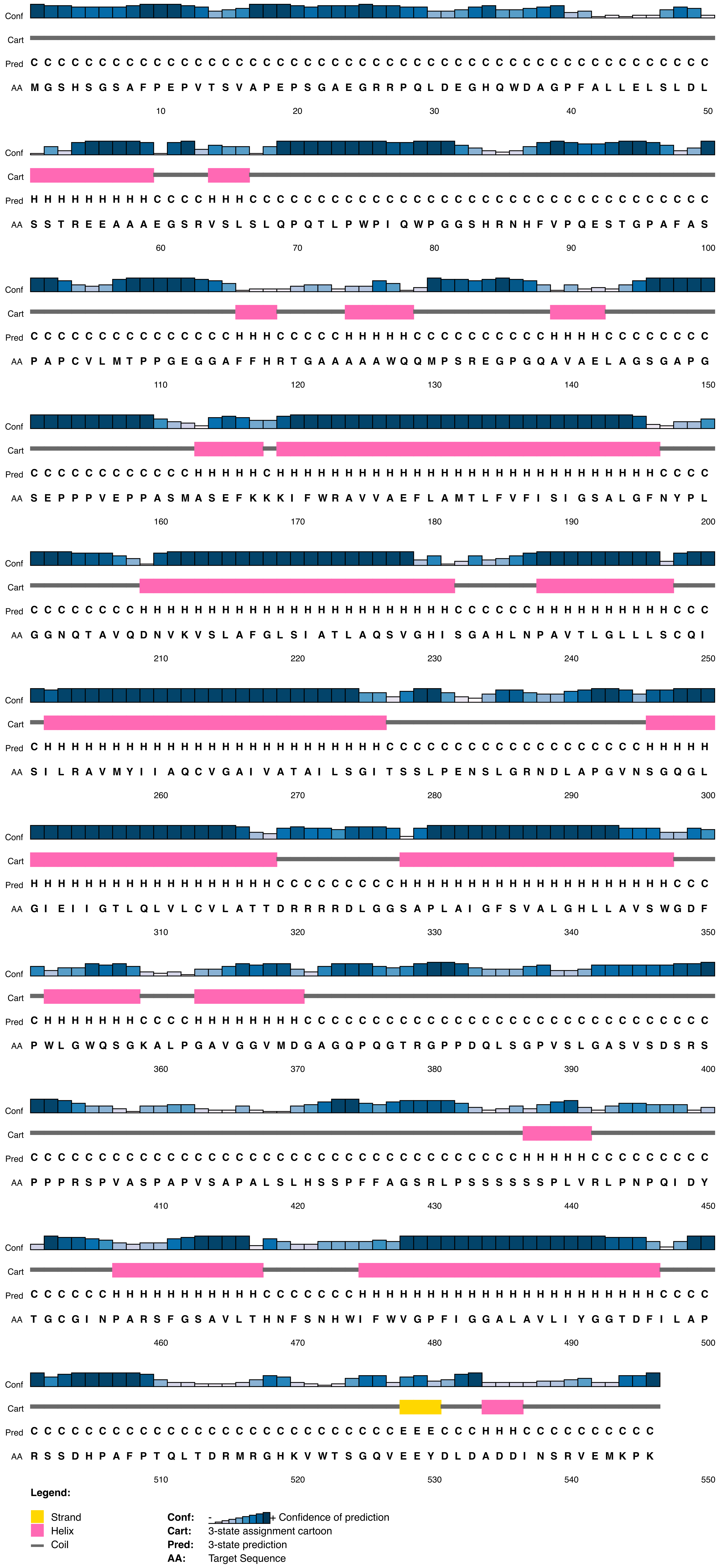


**Figure A4.10** – *Aqp1* secondary structure prediction for Anc 22. At the site under positive selection Anc 22 and its descendent *Oryctolagus cuniculus* both had a proline (see Figure 6 in the main text).

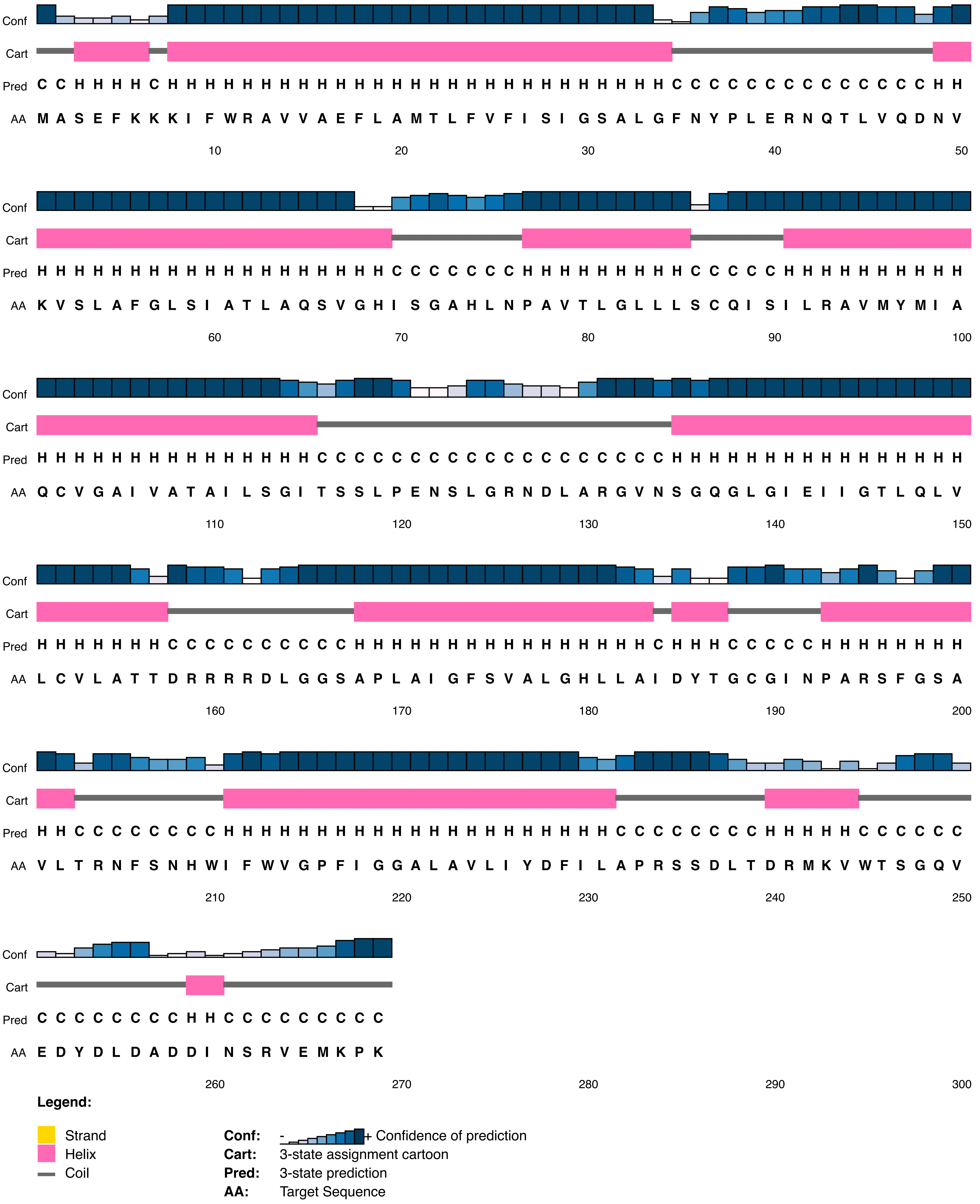


**Figure A4.11** – *Aqp1* secondary structure prediction for *Microtus ochrogaster*. At the site under positive selection, *M. ochrogaster* had an arginine and its ancestor, Anc8, had a proline (see Figure 6 in the main text).

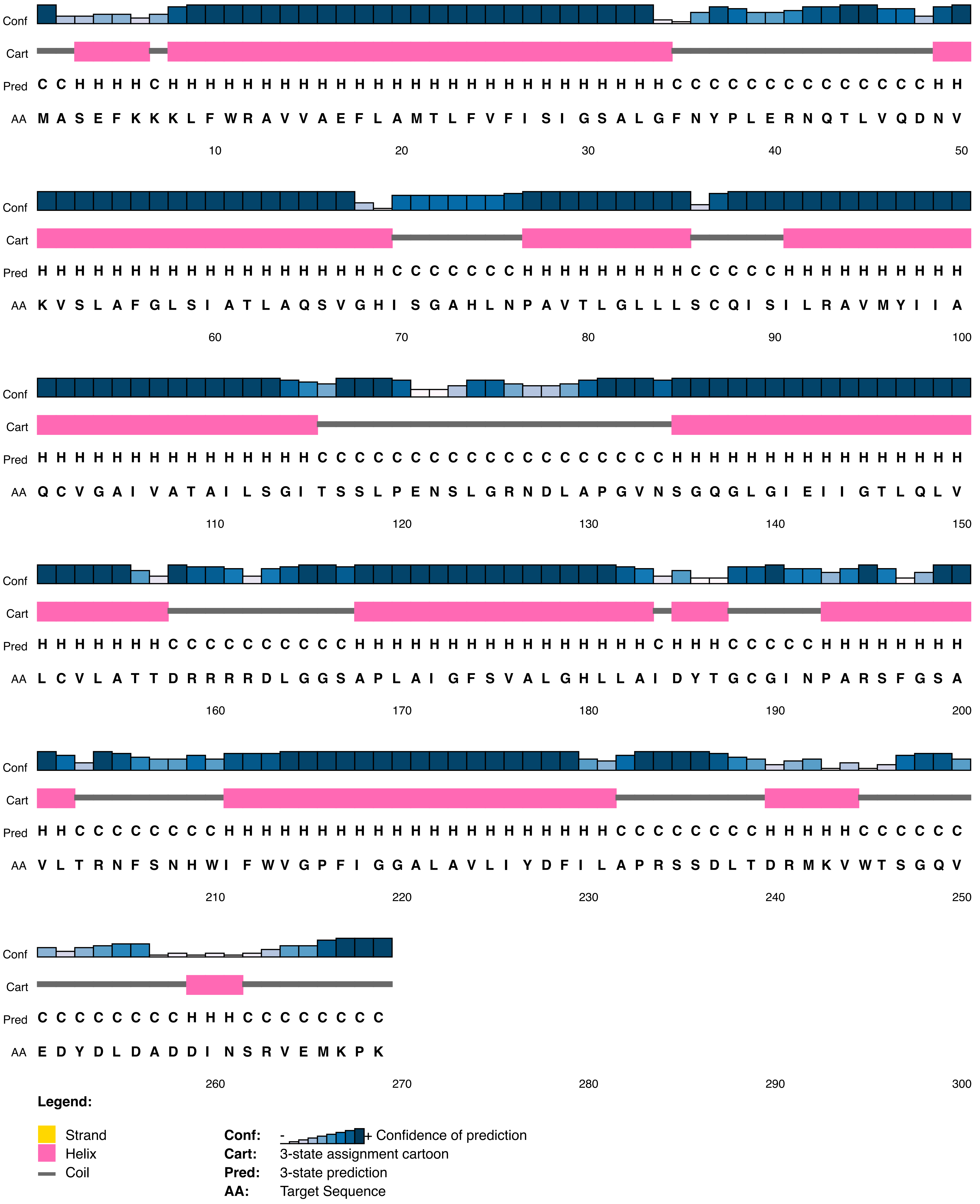


**Figure A4.12** – *Aqp1* secondary structure prediction for Anc8, the ancestor of *Microtus ochrogaster*. Anc8 had a proline at the site under positive selection which was subsituted for an arginine in *M. ochrogaster* (see Figure 6 in the main text).

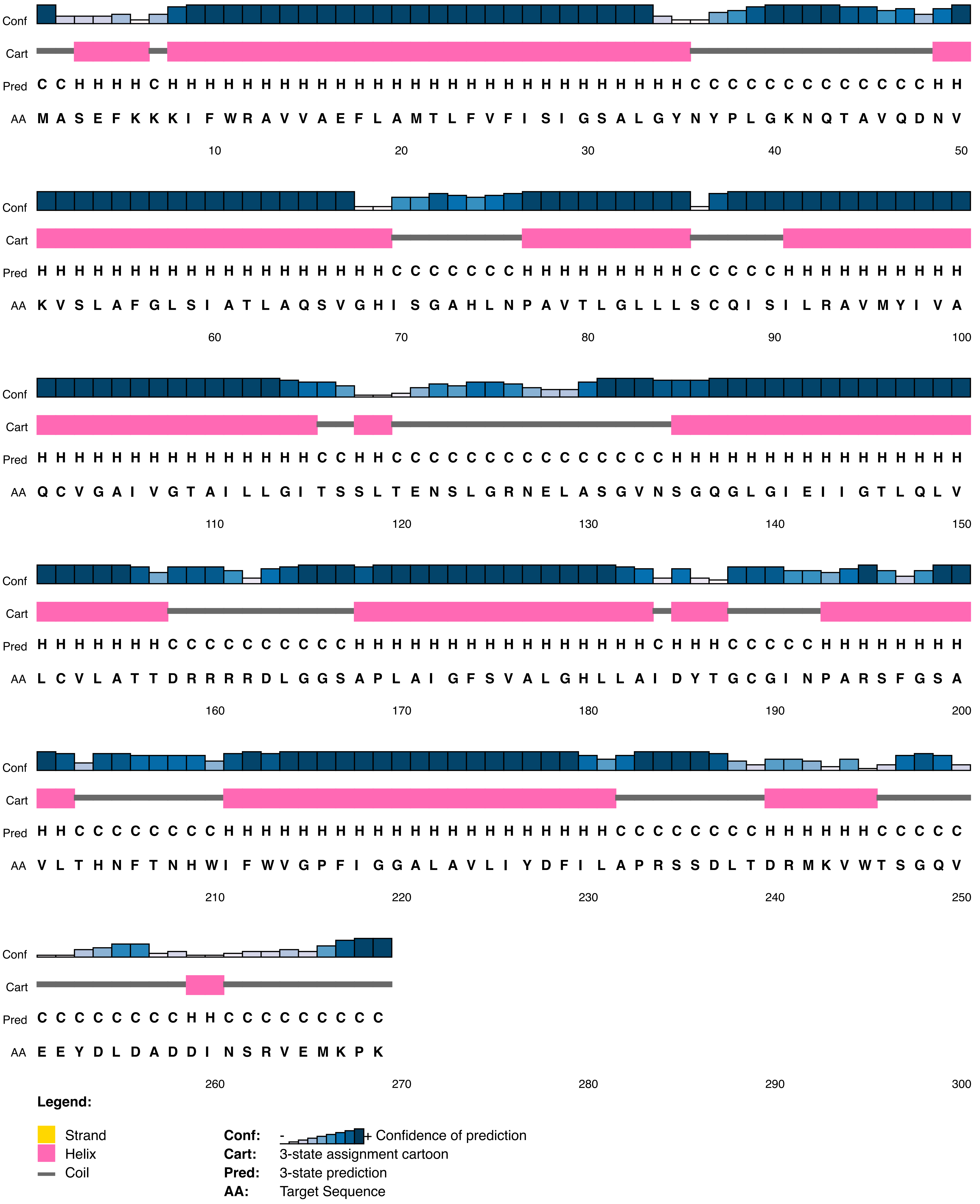


**Figure A4.13** – *Aqp1* secondary structure prediction for *Dipodomys ordii*. At the site under positive selection, *D. ordii* had an serine and its ancestor, Anc20, had a proline (see Figure 6 in the main text).

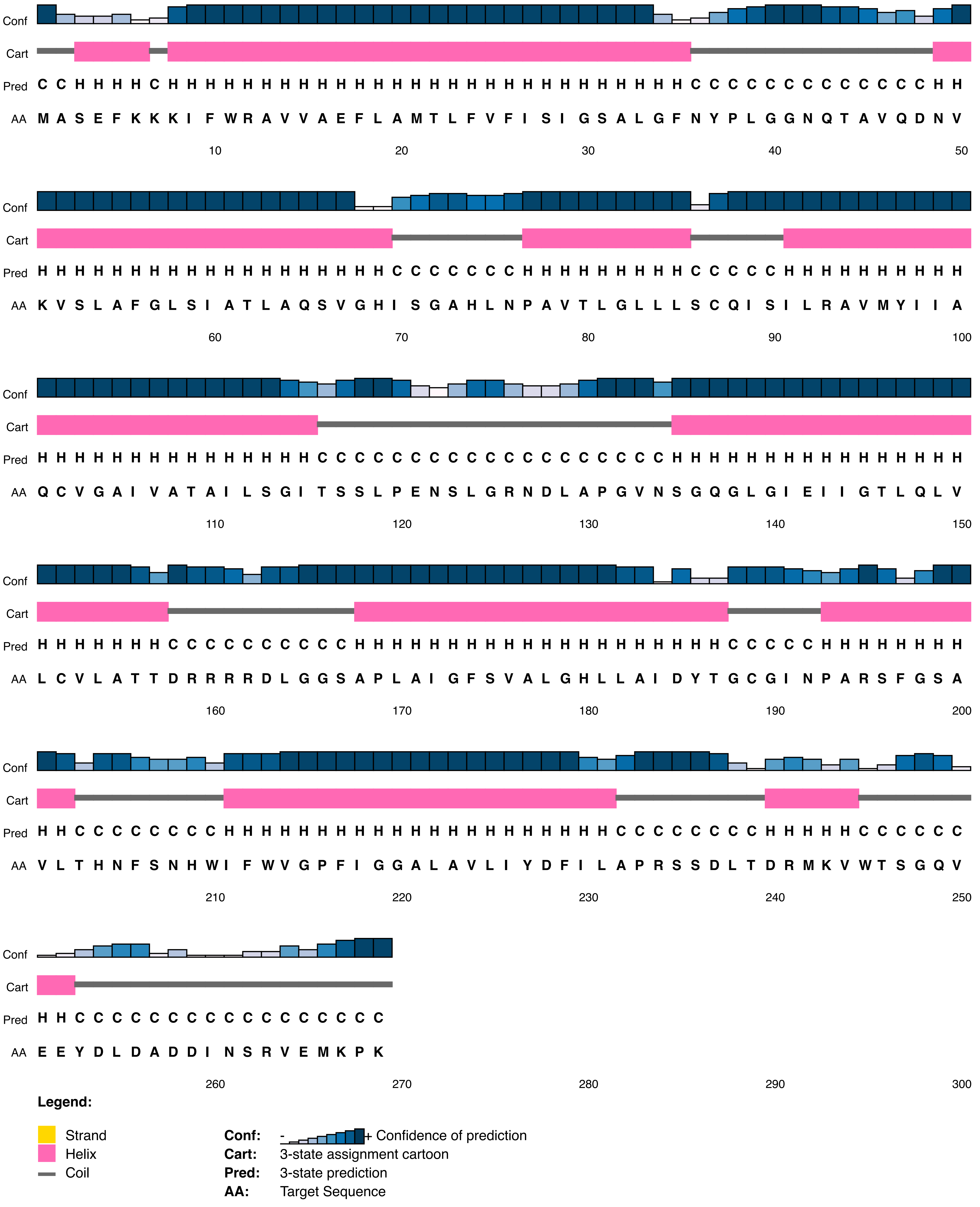


**Figure A4.14** – *Aqp1* secondary structure prediction for Anc20, the ancestor of *Dipodomys ordii*. Anc20 had a proline at the site under positive selection which was subsituted for an serine in *D. ordii* (see Figure 6 in the main text).

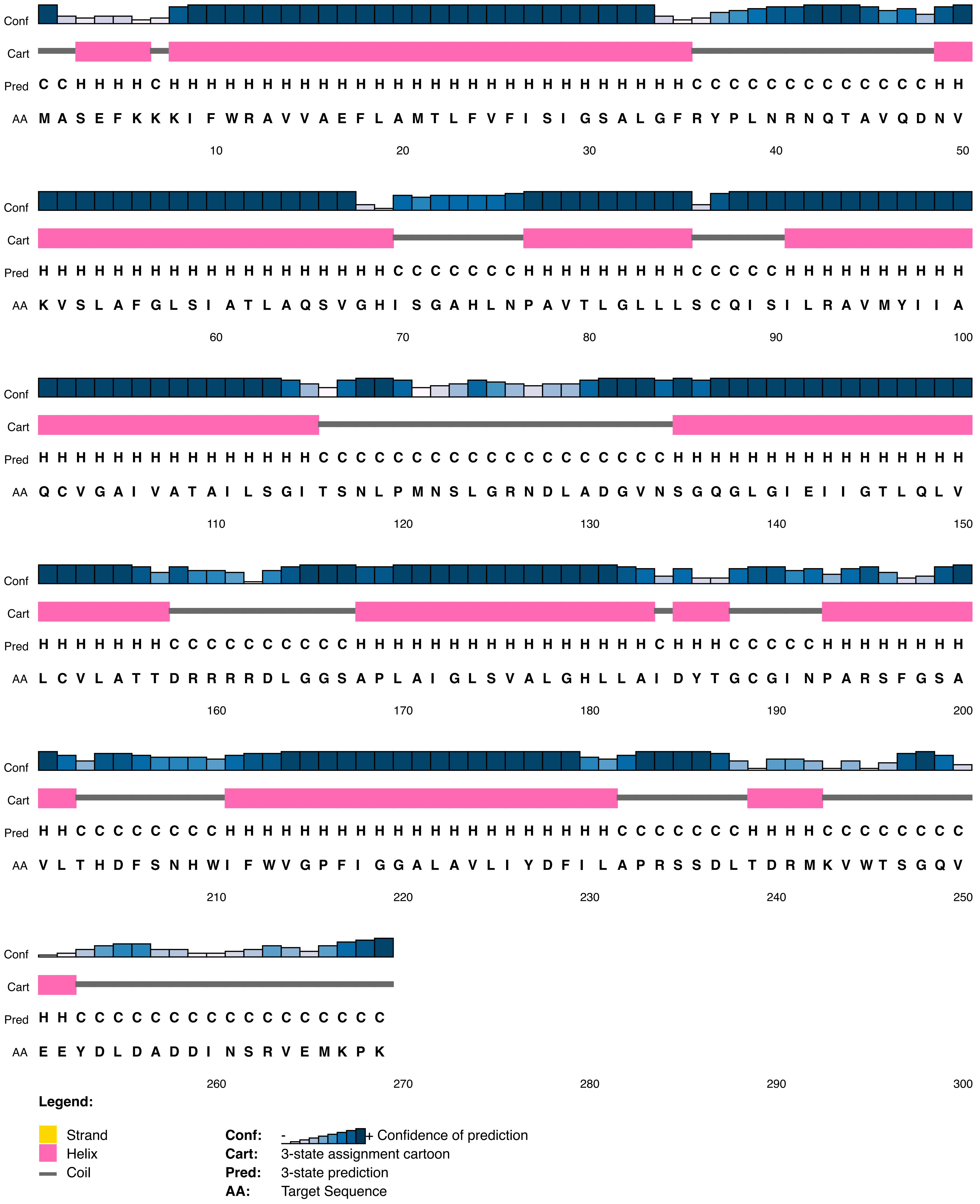


**Figure A4.15** – *Aqp1* secondary structure prediction for *Octodon degus*. At the site under positive selection, *O. degus* had an aspartic acid and its ancestor, Anc17, had a serine (see Figure 6 in the main text).

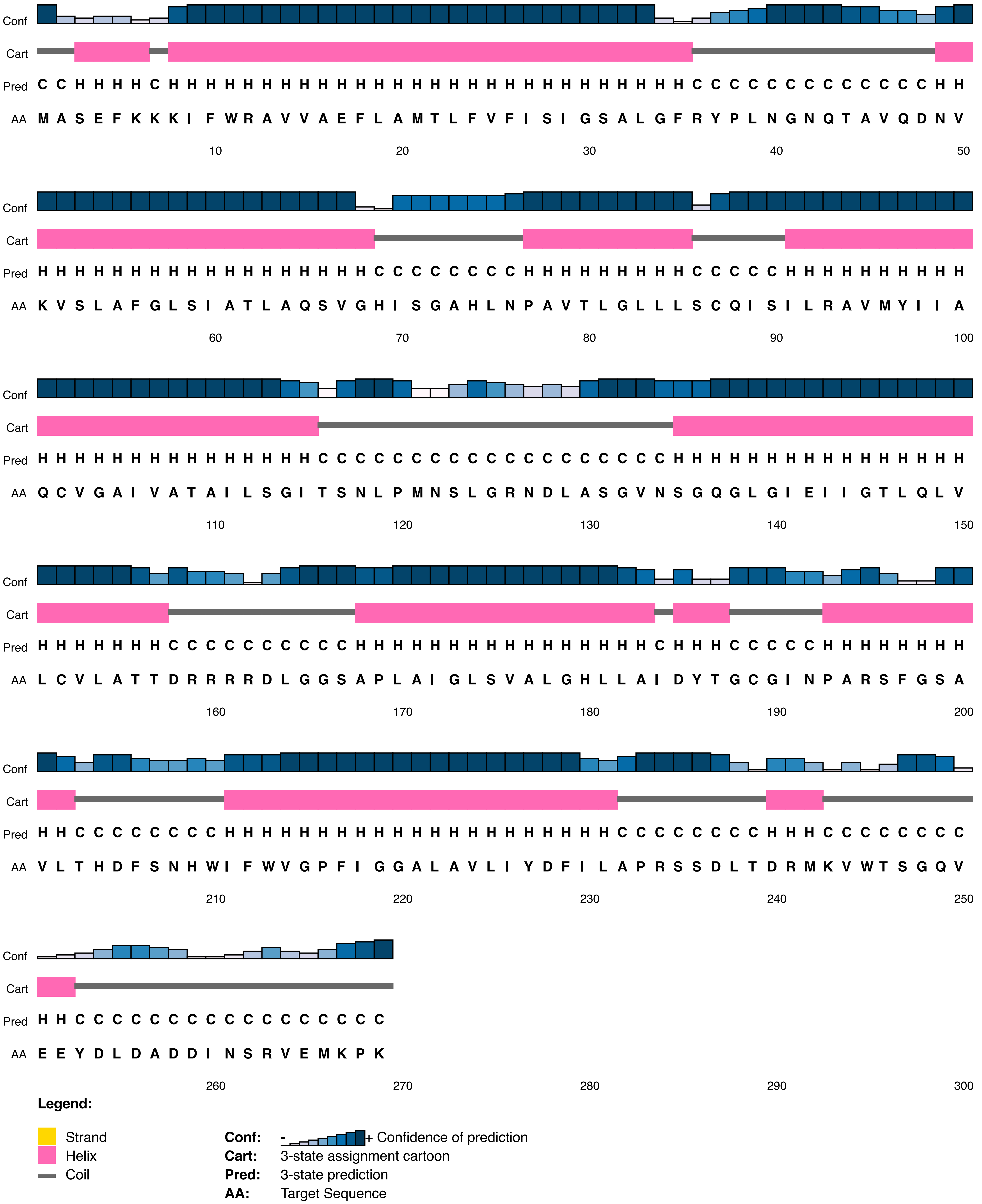


**Figure A4.16** – *Aqp1* secondary structure prediction for Anc17, the ancestor of *Octodon degus*. Anc17 had a serine at the site under positive selection which was subsituted for an aspartic acid in *O. degus* (see Figure 6 in the main text).

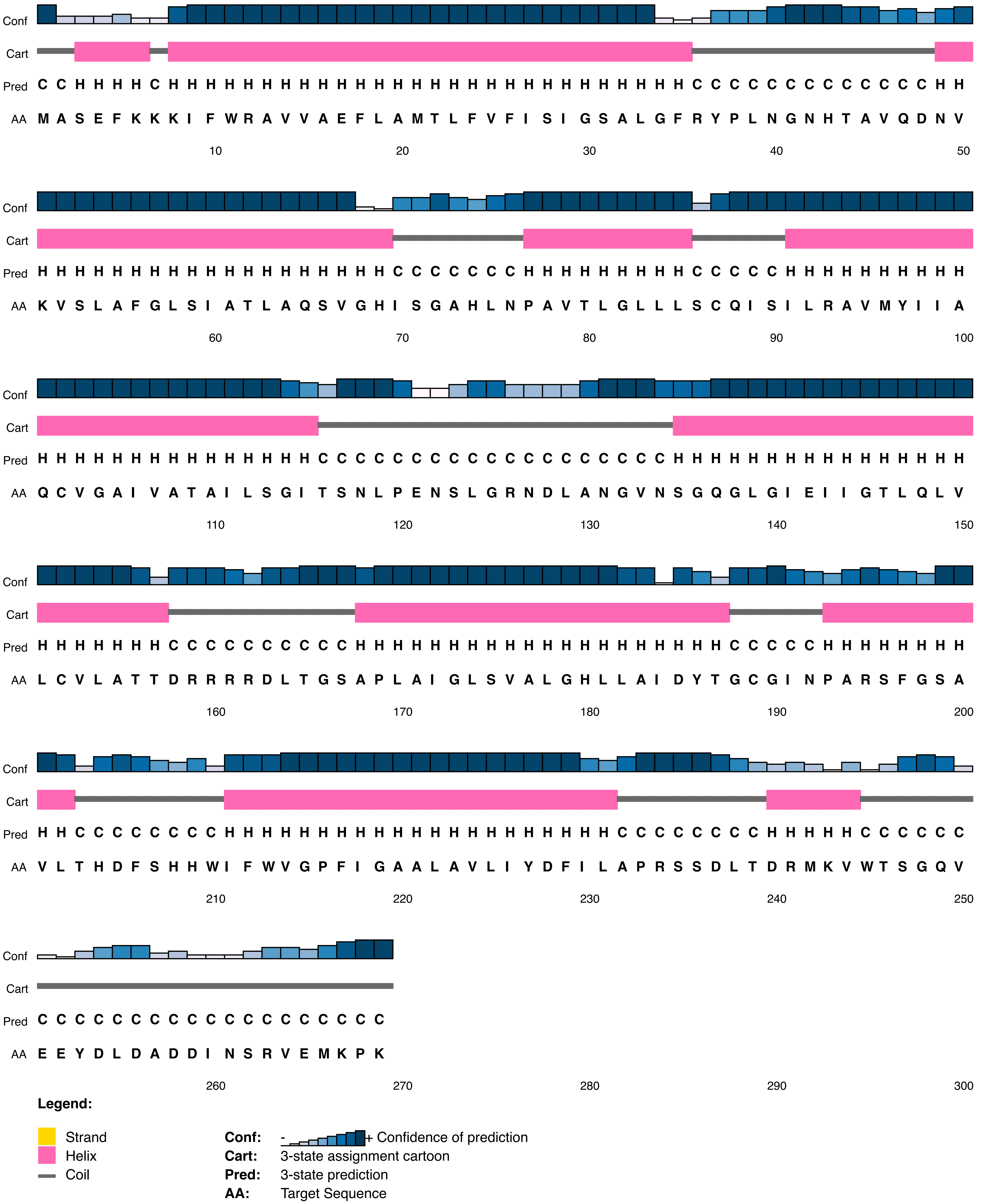


**Figure A4.17** – *Aqp1* secondary structure prediction for *Chinchilla lanigera*. At the site under positive selection, *C. lanigera* had an asparagine and its ancestor, Anc16, had a serine (see Figure 6 in the main text).

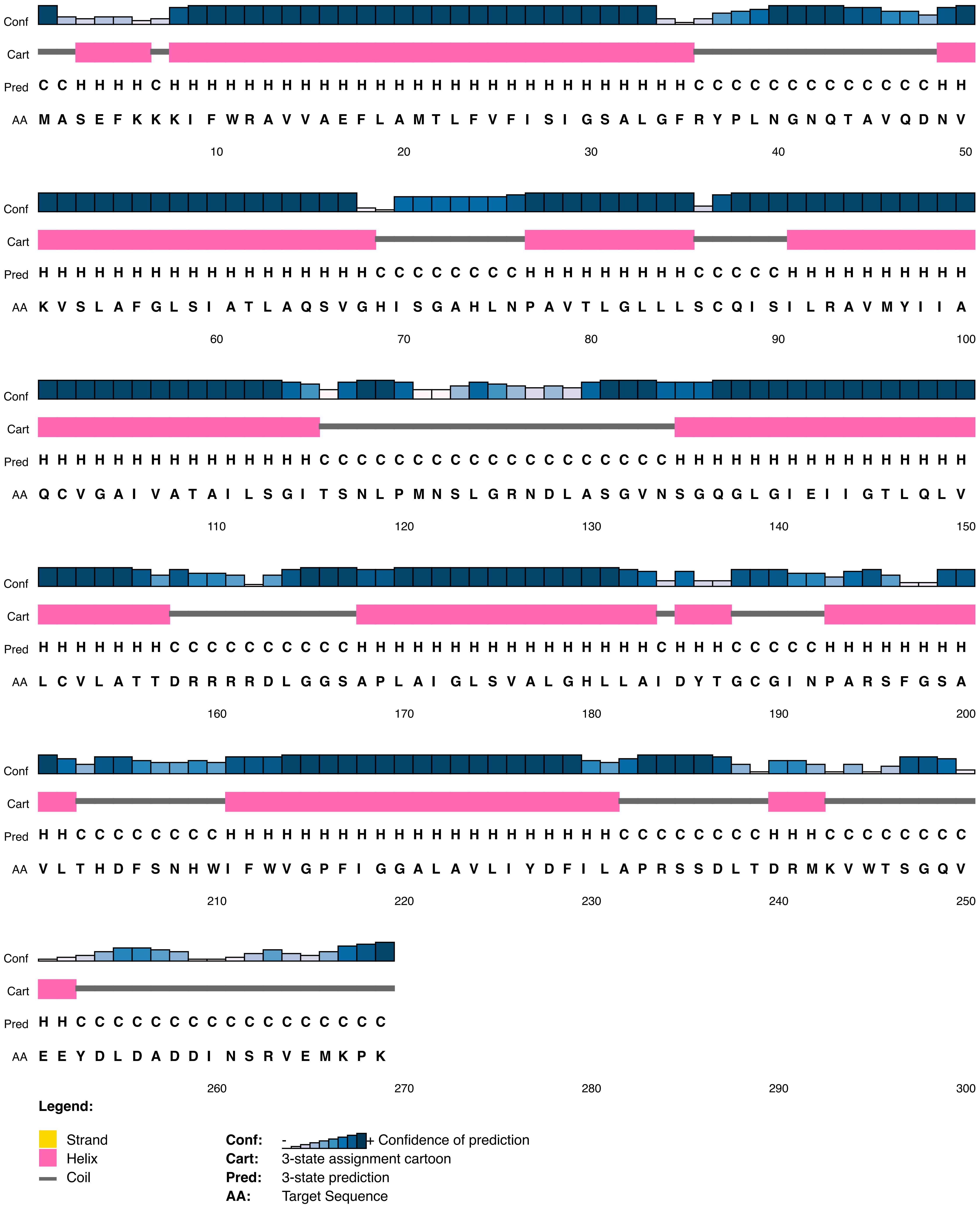


**Figure A4.18** – *Aqp1* secondary structure prediction for Anc16, the ancestor of *Chinchilla lanigera*. Anc16 had an serine at the site under positive selection which was subsituted for an asparagine in *C. lanigera* (see Figure 6 in the main text).

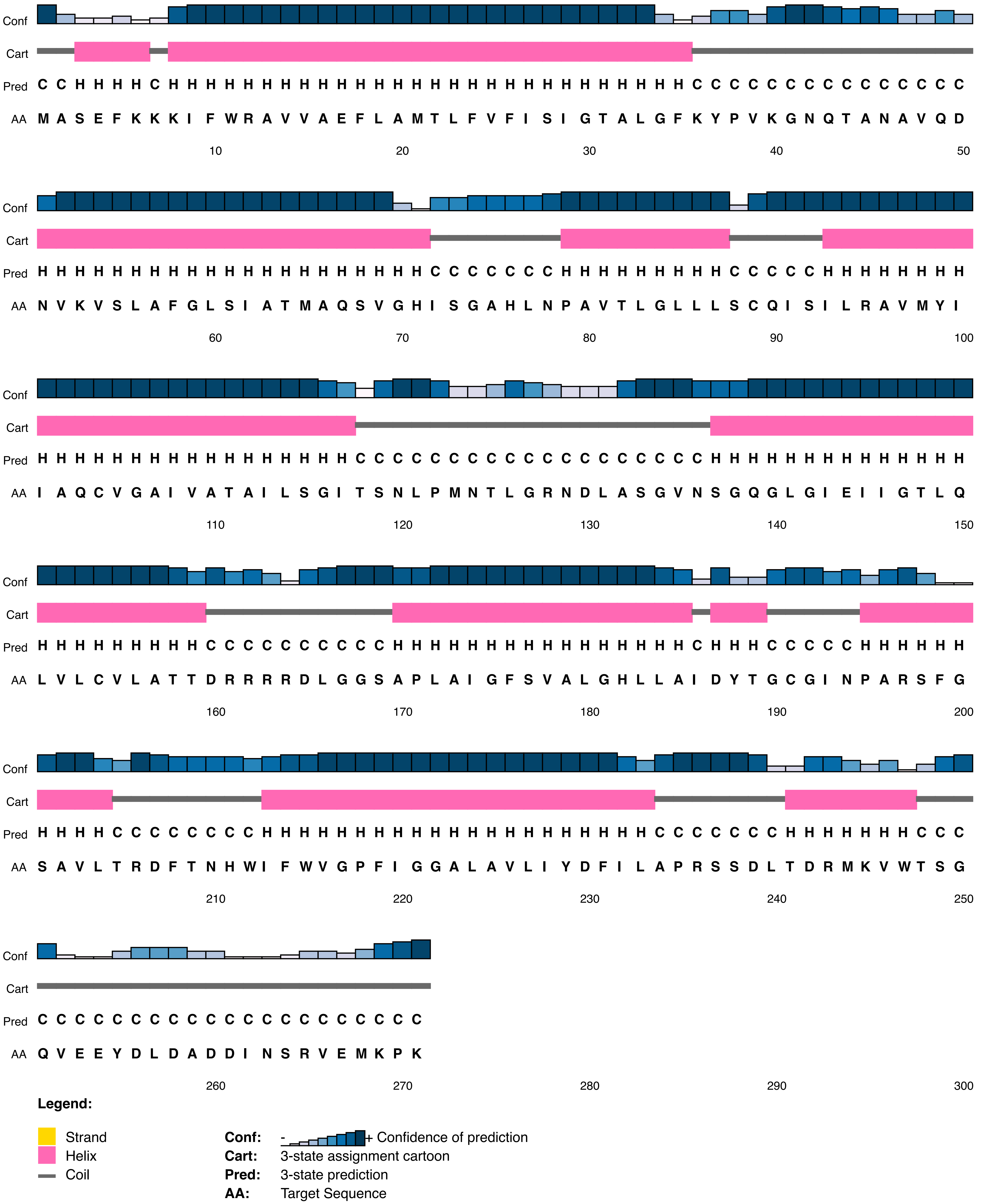


**Figure A4.19** – *Aqp1* secondary structure prediction for *Cavia aperea*. At the site under positive selection, *C. aperea* had a serine and its closest ancestor with a different amino acid, Anc18, had a proline (see Figure 6 in the main text).

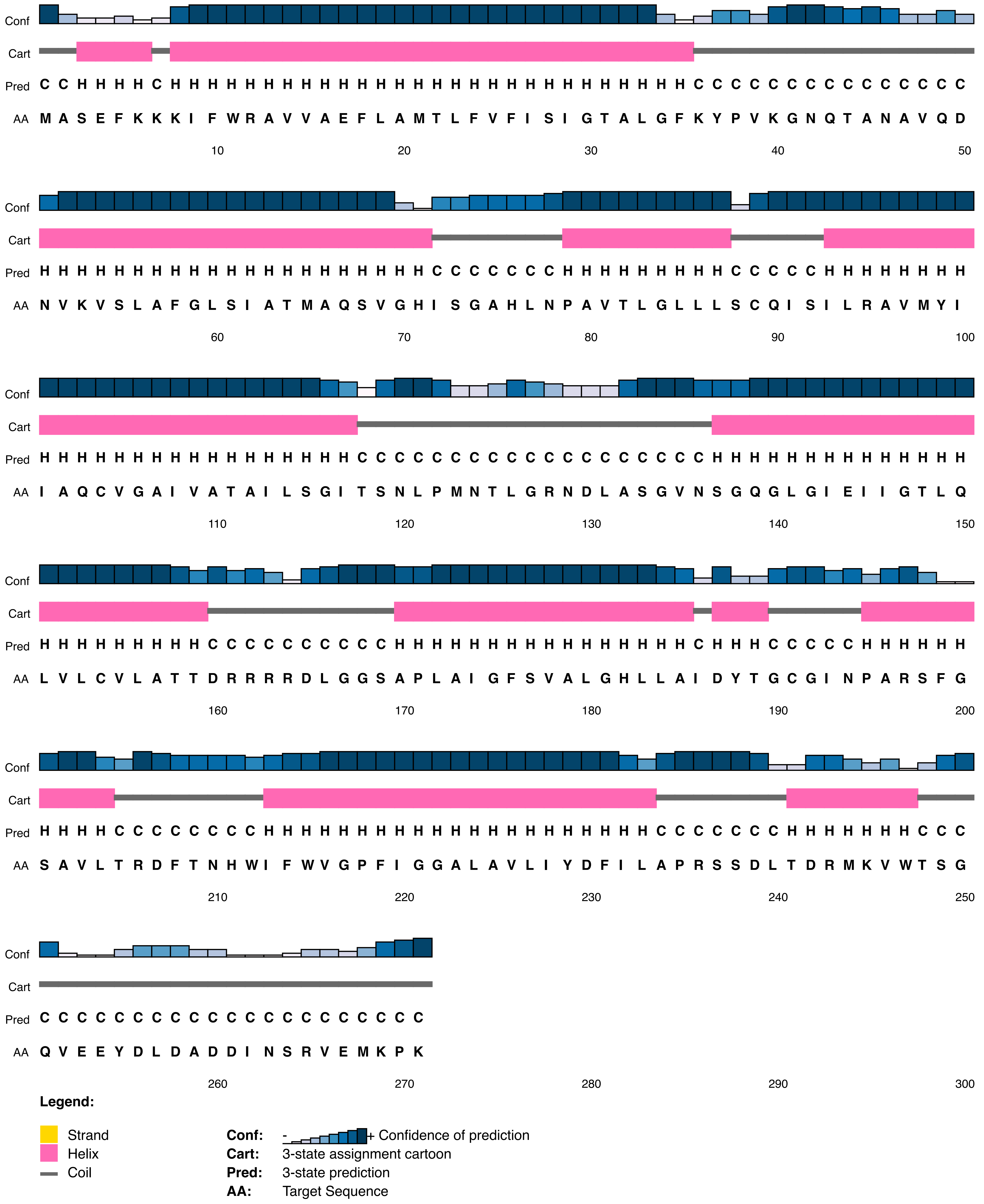


**Figure A4.20** – *Aqp1* secondary structure prediction for *Cavia porcellus*. At the site under positive selection, *C. porcellus* had a serine and its closest ancestor with a different amino acid, Anc18, had a proline (see Figure 6 in the main text).

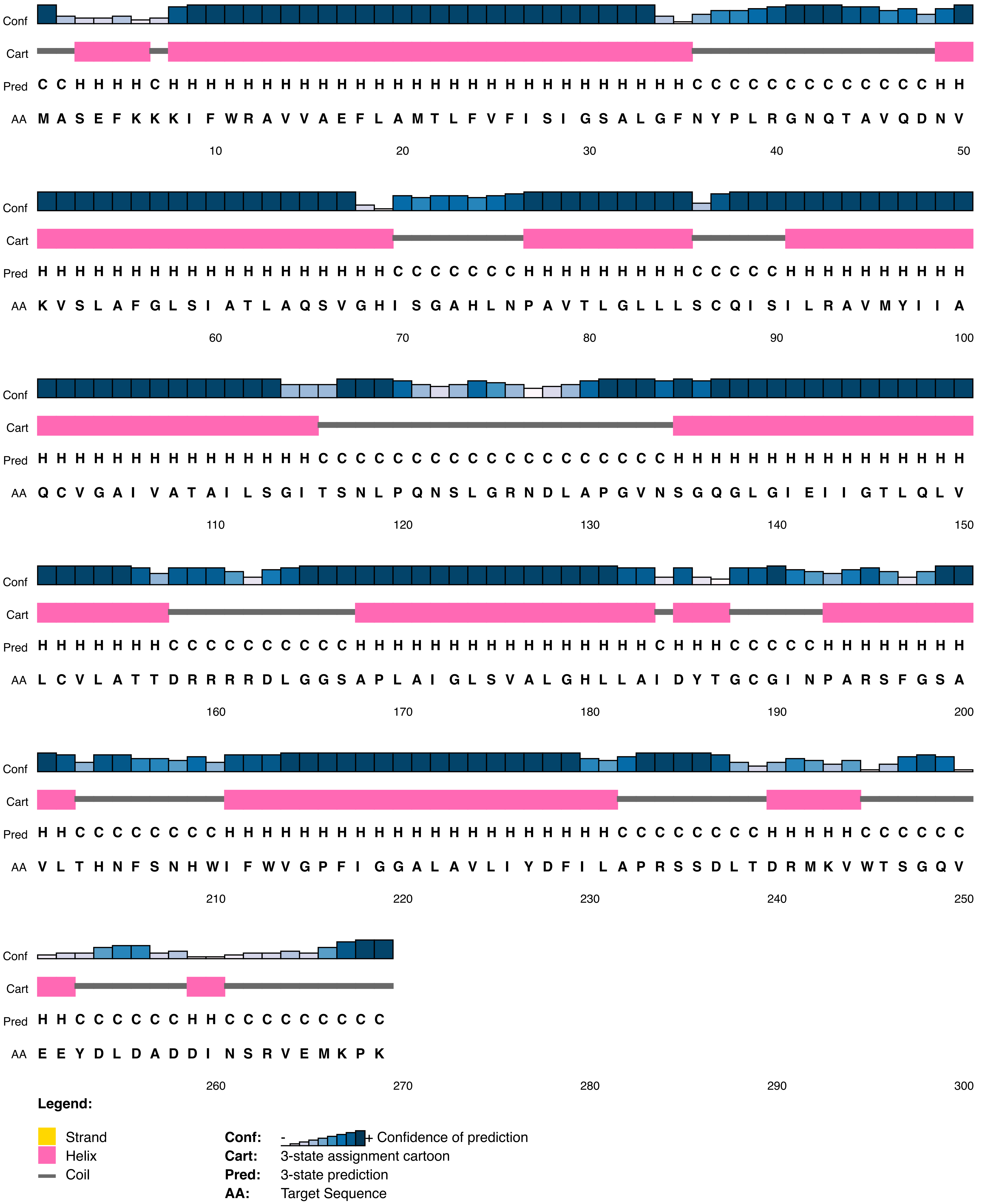


**Figure A4.21** – *Aqp1* secondary structure prediction for Anc18, the closest ancestor to *Cavia aperea* and *Cavia porcellus* with a different amino acid at the site under positive selection. Anc 18 had a proline at the site which was subsituted for a serine in both living species (see Figure 6 in the main text).
