## Additional file 6: Oligonucleotide primers for "Vole genomics links determinate and indeterminate growth of teeth"

Primer sequences used for qPCR analyses of expression in bank vole and prairie vole M1 at three embryonic days.

Bank vole (*Myodes glareolus*, rooted molars)
Aqp1_forward: GGGCATTGAGATCATCGGCA
Aqp1_reverse: CCAGTGTAGTCAATCGCCAG
Dspp_forward: AGGAACTCCAGCACAGAATGA
Dspp_reverse: TCGTCCCTCCTACGTCTGTT
GAPDH_forward: GTGGGCAAAGTCATCCCAGA
GAPDH_reverse: GTGTAGCCCTTGATGCCCTT

Prairie vole (*Microtus ochrogaster*, unrooted molars)
Aqp1_forward: GCTCCTGCTCAGTTGTCAGAT
Aqp1_reverse: CACACCTCGAGCCAGGTCATT
Dspp_forward: GGAACTCCAGCACAGGATGA
Dspp_reverse: TCGTCCCTCCTACGTCTGTT
GAPDH_forward: TGGAGACAGCCGCTTCTTTT
GAPDH_reverse: GCGTCCAATACGGCCAAATC
